# Cdc42 reactivation at growth sites is regulated by local cell-cycle-dependent loss of its GAP Rga4

**DOI:** 10.1101/2020.08.14.238121

**Authors:** Julie Rich-Robinson, Afton Russell, Eleanor Mancini, Maitreyi Das

## Abstract

In fission yeast, polarized cell growth stops during division and resumes after cell separation. We uncoupled these sequential events by delaying cytokinesis with a temporary Latrunculin A treatment. Mitotic cells recovering from treatment initiate end growth without cell separation, displaying a **p**ola**r e**longation **s**ans **s**eparation (PrESS) phenotype. PrESS cell ends reactivate Cdc42, a major regulator of polarized growth, before cell separation, but at a fixed time after anaphase B. A candidate screen implicates Rga4, a negative regulator of Cdc42, in this process. We show that Rga4 appears punctate at the cell sides during G2, but is diffuse during mitosis, extending to the ends. While the Morphogenesis Orb6 (MOR) pathway is known to promote cell separation and growth by activating protein synthesis, we find that for polarized growth, removal of Rga4 from the ends is also necessary. Therefore, we propose that growth resumes after division once the MOR pathway is activated and the ends lose Rga4 in a cell-cycle-dependent manner.

## Introduction

Most eukaryotic cells undergo polarized growth to achieve specific shapes that enable them to interact with their environment in a biologically productive manner. Polarized growth requires polarization of growth-promoting signaling networks, actin organization, and membrane trafficking (Etienne-Manneville, 2004; Nance and Zallen, 2011; Ridley, 2006). Polarization is also essential for cytokinesis, the final step in cell division (Albertson et al., 2005; Echard, 2008; Hercyk et al., 2019a; Wang et al., 2016). Since polarized growth and cytokinesis share the same polarization apparatus, they must be temporally distinct events. Accordingly, careful regulation ensures that the cell can alternate between successful growth and division. However, it is unclear how this is regulated at the molecular level. The major drivers of polarization are highly conserved among eukaryotes (Etienne-Manneville, 2004; Johnson, 1999). For this reason, we use the unicellular fission yeast *Schizosaccharomyces pombe* to establish basic principles about how cells spatiotemporally regulate their polarization machinery.

In fission yeast, cytokinesis involves the formation of an actomyosin ring, which constricts along with an ingressing membrane barrier and septum to physically separate two daughter cells (Cheffings et al., 2016; Pollard, 2010; Pollard, 2014). The septum undergoes maturation and is eventually digested by glucanases, leading to cell separation (Cortes et al., 2016; Garcia Cortes et al., 2016; Sipiczki, 2007). Immediately after cell separation, polarized growth initiates from the old ends, which existed in the previous generation (Mitchison and Nurse, 1985). Once the cells attain a certain size, the new ends, generated as a result of cell division, start to grow by a process called new end take-off (NETO). The Rho GTPase Cdc42 is a major regulator of cell growth and polarity, and is highly conserved among eukaryotes (Etienne-Manneville, 2004; Johnson, 1999). Recently, we have shown that Cdc42 also plays a role in cytokinesis (Hercyk and Das, 2019a; Onwubiko et al., 2020; Wei et al., 2016). Regulation of Cdc42 determines when and where polarization occurs (Das et al., 2012; Das et al., 2009; Etienne-Manneville, 2004; Howell et al., 2012). In *S. pombe*, Cdc42 is activated by two Guanine nucleotide exchange factors (GEFs), Scd1 and Gef1 (Chang et al., 1994; Coll et al., 2003), and is inactivated by three GTPase-activating proteins (GAPs), Rga4, Rga6, and Rga3 (Das et al., 2007; Gallo Castro and Martin, 2018; Revilla-Guarinos et al., 2016; Tatebe et al., 2008). Of these, Rga4 appears to be the primary GAP, while Rga6 and Rga3 appear to play minor roles in polarity and cell pairing during sexual reproduction, respectively.

The ends of dividing cells inactivate Cdc42 during mitosis, and polarized growth consequently stops (Hercyk and Das, 2019b). Once two daughter cells have been generated via cytokinesis at the end of cell division, Cdc42 is reactivated at the old ends and polarized growth resumes (Wei et al., 2016). The bio-probe CRIB-3xGFP specifically binds active Cdc42, and thus is used to identify sites of Cdc42 activation (Tatebe et al., 2008). Using CRIB-3xGFP, it has been shown that Cdc42 is activated at growing cell ends during interphase (Das et al., 2012; Tatebe et al., 2005). Once the cell enters mitosis, active Cdc42 disappears from the cell ends and appears at the division site (Wei et al., 2016). After cell separation, Cdc42 is not immediately activated at a daughter cell’s new end, which was created as a result of cell separation. Instead, Cdc42 is first activated at the old end, which is the farthest point from the site of cell division. It is unclear how Cdc42 activation transitions from the division site to the old ends after cell separation.

While much is understood about *how* polarized growth occurs, little is known about the regulation of *when* polarized growth occurs. It has been shown that cell-cycle-dependent signaling pathways temporally segregate cell division and growth phases (Ray et al., 2010; Simanis, 2015). During interphase, the Morphogenesis Orb6 (MOR) pathway promotes cell growth (Nunez et al., 2016; Ray et al., 2010). During cell division, the septation initiation network (SIN) is activated, allowing septum formation (Johnson et al., 2012; Simanis, 2015). Activation of the SIN leads to inactivation of the MOR pathway, thus inhibiting cell growth during septum closure (Ray et al., 2010).

Subsequent SIN inactivation allows MOR pathway activation, which leads to cell separation and growth (Gupta et al., 2014; Gupta et al., 2013). This crosstalk between the SIN and MOR signaling pathways thus enforces temporal separation of cytokinesis and cell growth.

The MOR pathway promotes cell separation and polarized growth activation, which occur sequentially. It is unclear how the MOR pathway promotes these distinct, sequential cellular processes. The MOR pathway involves activation of the Orb6/NDR kinase, which phosphorylates the exoribonuclease Sts5 (Nunez et al., 2016). Phosphorylation of Sts5 prevents its incorporation into processing bodies (P-bodies), where mRNA is stored or degraded (Nunez et al., 2016). Orb6 activity thus prevents Sts5-dependent degradation of mRNA in the P-bodies, leading to increased protein synthesis. Among the mRNAs spared from degradation are those that encode the glucanases Eng1 and Agn1, and the Ras1 GEF Efc25 (Chen et al., 2019; Nunez et al., 2016). While Eng1 and Agn1 promote septum digestion and cell separation, Efc25- mediated Ras1 activation promotes the localization of the Cdc42 GEF Scd1 to cell ends (Garcia et al., 2005; Lamas et al., 2020a; Martin-Cuadrado et al., 2003; Papadaki et al., 2002). In this way, Orb6 activity promotes cell separation and polarized cell growth. However, constitutively activating the MOR pathway does not lead to constitutive cell growth, but rather to premature cell separation that results in cell lysis (Gupta et al., 2014). This suggests that, once the MOR pathway is activated, it immediately promotes both cell separation and growth initiation. Thus, the nature of the regulation that allows cell separation and growth initiation to occur sequentially is unclear, and is the focus of this study.

Here we investigate how growth is reactivated at the cell ends after cell division. After artificially delaying cytokinesis, we observe that the timing of growth activation at the ends is not influenced by the completion of cytokinesis and consequent cell separation. In keeping with this, we find that Cdc42 activity at the cell ends resumes at a fixed time after completion of anaphase B and is independent of cell separation. Our candidate screen for regulators involved in reactivation of Cdc42 at the cell ends identified the Cdc42 GAP Rga4. Rga4 has previously been shown to localize to the cell sides, where it prevents Cdc42 activation (Das et al., 2007; Tatebe et al., 2008). We find that the localization pattern of Rga4 along the cortex changes in a cell-cycle-dependent manner. During interphase, Rga4 appears punctate along the cortex and is excluded from cell ends. During mitosis, Rga4 appears diffuse and localizes all along the cortex, including the cell ends. We propose that Rga4 localization at the cell ends during mitosis blocks Cdc42 activation at these ends. Upon completion of mitosis, Rga4 is lost from the ends and Cdc42 activity resumes. This suggests that once mitosis completes, growth activation at the cell ends occurs once Rga4 is lost from these ends. MOR pathway activation promotes both cell separation and end growth, which normally occur sequentially. Our data suggest that polarized growth activation occurs only once Rga4 is lost from growth sites, allowing these events to occur in sequence. In keeping with our findings, we show that, in mutants constitutively activating the MOR pathway, loss of *rga4* leads to enhanced polarized cell growth. These data suggest that cell-cycle- dependent regulation of Rga4 paired with MOR pathway activation allow cell separation and polarized cell growth to occur sequentially.

## Results

### Cytokinetic delay results in growth resumption without cell separation

Since initiation of polarized growth at cell ends only occurs after cell separation, we asked if the timing of polarized growth initiation is impacted by cytokinesis. To test this, we used Latrunculin A (LatA) to induce a cytokinetic delay. LatA prevents the polymerization of actin, leading to loss of dynamic actin-based structures (Spector et al., 1983). Phalloidin staining shows loss of actomyosin rings upon LatA treatment (Supplemental Figure S1A). LatA treatment disrupts the actomyosin ring, and upon washout, a new actomyosin ring assembles and cytokinesis resumes, albeit it is delayed (Supplemental Figure S1B) (Swulius et al., 2018). We treated an asynchronous population of wild type cells with 10µM LatA for 30 minutes and then washed it out.

Cells typically take about 45-60 minutes after LatA washout to recover before they display growth. After recovery, we observed that some cells – about 9% – show a unique phenotype in which they initiate growth at their ends but do not undergo cell separation (Figure 1A, B; Supplemental Movie S1). We analyzed the timing of end growth initiation with reference to completion of septum closure in these cells. We monitored cells using bright-field imaging over time to detect septum closure (Supplemental Figure S1C). In untreated wild type cells, growth at the ends initiates on average 50 minutes after septum closure. However, in cells recovering from LatA treatment, end growth initiates on average 6 minutes after septum closure (Figure 1D). In some cells, growth initiates even before the completion of septum closure. We observe similar results when growth resumption is timed from the completion of ring closure, as depicted by Rlc1-tdTomato (Supplemental Figure S1D). Although these cells fail to separate, they enter the next cell cycle and undergo cell division. The septum formed in the second generation separates normally, suggesting that the LatA treatment only impacts the cell cycle in which the cell was treated (Figure 1A, arrow). We call this phenotype the **<u>p</u>**ola**<u>r e</u>**longation **<u>s</u>**ans **<u>s</u>**eparation, or PrESS, phenotype. We also observe that the old ends of PrESS cells grow up to three times as fast as old ends in wild type cells before new end take-off (Supplemental Figure S1E). The PrESS phenotype suggests that growth initiation after mitosis is not timed by cytokinesis.

**Figure 1.**
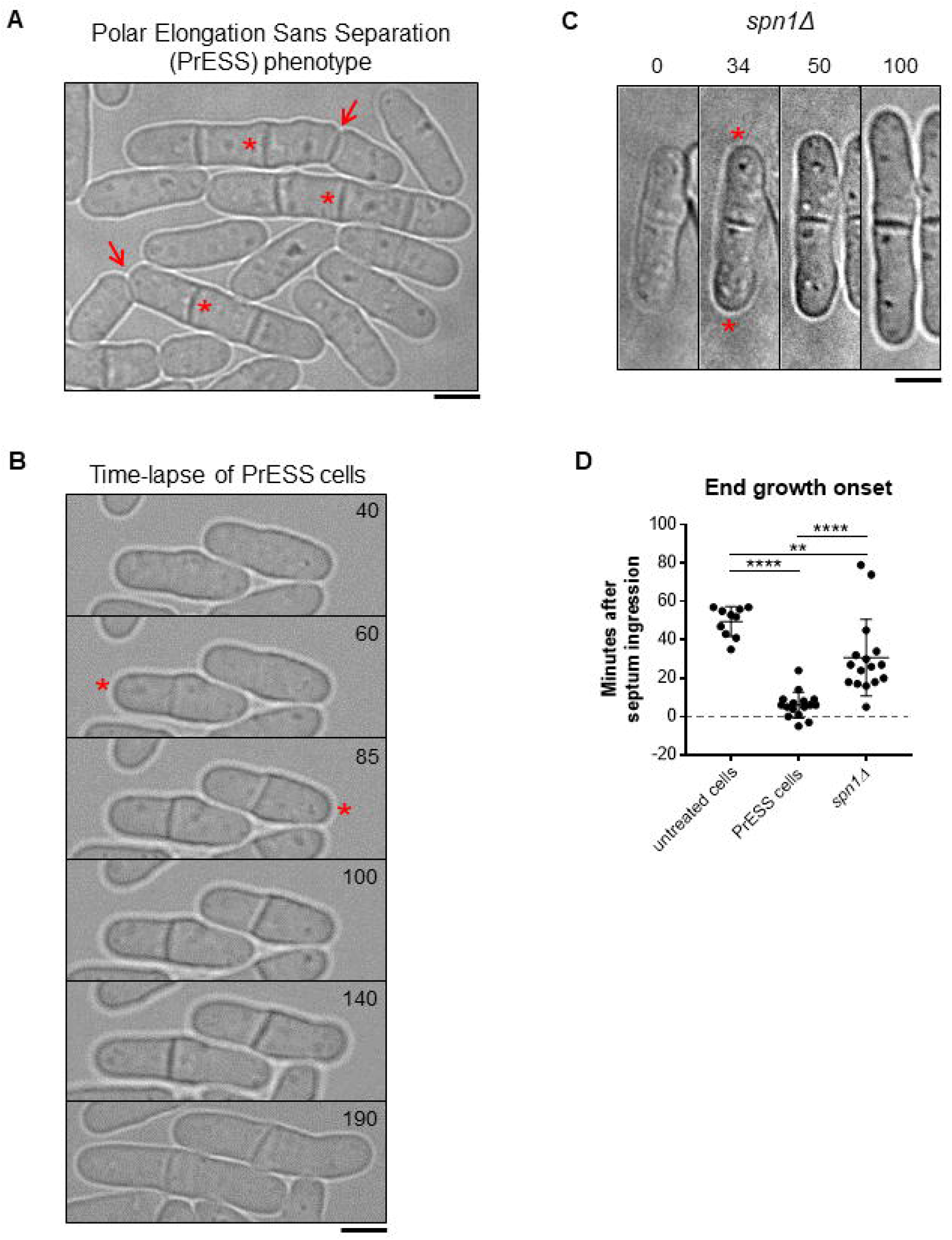
Cytokinetic delay uncouples end growth resumption from cell separation. **A)** An asynchronous population of wild type cells was treated with 10µM Latrunculin A (LatA) for 30 minutes, and was then washed and allowed to recover. A subset of cells consistently shows a phenotype wherein they form a septum as they resume growth and fail to undergo cell separation (asterisk next to original septum). These cells grow, and the septa formed in the next cell cycle separate normally (arrows). We call this the **<u>p</u>**ola**<u>r e</u>**longation **<u>s</u>**ans **<u>s</u>**eparation, or PrESS, phenotype. **B)** Time-lapse of two wild type cells after LatA washout, showing the PrESS phenotype. Growth resumed in the left cell at 60mins, and in the right cell at 85mins after washout (asterisks). **C)** Time-lapse of growth after division in an *spn1Δ* cell. Asterisks denote growth initiation, which occurs after septum formation. **D)** PrESS cells resume growth significantly earlier in relation to septum closure than untreated or *spn1Δ* cells (Student’s t-test, p < 0.0001; n ≥ 10 cells). Timestamps in B and C refer to time since completion of septum closure. Scale bars, 5μm.

PrESS cells visually resemble septin mutants, in which cell separation defects have been previously documented (Figure 1C) (An et al., 2004; Berlin et al., 2003; Martin-Cuadrado et al., 2005; Tasto et al., 2003). However, while PrESS cells resume growth as the septum forms, it was unclear if the *spn1Δ* mutants behave similarly or whether they only resume growth after cell separation failure. To test this, we analyzed the timing of end growth resumption in *spn1Δ* mutants compared to the time of septum closure, as we did for PrESS cells. We find that in *spn1Δ* cells, the ends resume growth on average about 31 minutes after septum closure in cells which fail to separate (Figure 1D). This suggests that these cells fail to separate, and then eventually growth resumes at the proper time. In contrast, in PrESS cells, cytokinesis and growth are uncoupled such that growth resumes even before cell separation is possible.

### Cdc42 activation at cell ends is cell-cycle-dependent

Only a subset of asynchronous cells exhibits the PrESS phenotype upon recovery from LatA treatment. We asked if this is because the PrESS phenotype only occurs in cells that are in a certain cell cycle stage. To address this, we imaged cells at various cell-cycle stages before, during, and after LatA treatment. We accomplished this by affixing the cells to the imaging dish using lectin (Tay et al., 2018), and performing LatA treatment and washout within the dish itself. Cells in lectin-coated dishes tend to grow slower than normal, but do not show any significant growth defects. We found that mitotic cells treated with LatA show the PrESS phenotype after recovery (Figure 2Aii, Supplemental Figure S2A, Supplemental Movie S2), and that cells treated during interphase or G1/S separate normally (Figure 2Ai, iii, Supplemental Figure S2A). When treated with LatA, cells in anaphase continue to proceed through mitosis, as detected with the spindle pole body marker Sad1-mCherry. However, the actomyosin ring disassembles, as marked by the type 2 myosin light chain Rlc1-tdTomato. After LatA washout, a new actomyosin ring assembles and then proceeds to constrict along with septum ingression (Figure 2Aii, arrowhead; Supplemental Figure S1B and C). However, instead of undergoing septum digestion and cell separation, these cells grow from the ends, thus displaying the PrESS phenotype. These observations show that the PrESS phenotype arises from cells that are in mitosis at the time of LatA treatment.

To confirm this, we synchronized *cdc25-22* cells by shifting them to restrictive temperature and arresting them in late G2 (Fantes and Nurse, 1978; Tormos-Perez et al., 2016). Upon shifting back to permissive temperature, the synchronized cells enter mitosis. We took samples of cells released from arrest at regular time intervals. The cell-cycle stage of these cells was determined by Phalloidin and DAPI staining after formaldehyde fixation of a fraction of the samples. Cells in G2 show a single nucleus with actin mainly at the cell ends. Cells in mitosis show a dividing nucleus with an actin ring at the cell middle (Supplemental Figure S2B). In G1/S, the cell middle shows a septum instead of an actin ring, with a single nucleus in each sister cell. The remaining fractions of the samples were treated with LatA like before and allowed to recover. We found that cells treated with LatA during late G2 went on to septate and divide normally (Supplemental Figure S2B). Cells released from restrictive temperature for 40 minutes were in mitosis at the time of LatA treatment. These cells almost exclusively displayed the PrESS phenotype during recovery. Cells released from restrictive temperature for 80 minutes were septated (in G1/S) at the time of LatA treatment, as evidenced by the divided nuclei and the lack of an actomyosin ring. These cells successfully separated after recovery and did not display the PrESS phenotype. To confirm that cells treated during mitosis were indeed able to grow from their ends, we looked for the presence of active Cdc42 at those ends. To do this, we treated *cdc25-22* mutants expressing CRIB-3xGFP, a probe for active Cdc42, in the same way as above. As expected, cells treated in G2 or G1/S underwent cell separation 2 hours after LatA washout and did not display the PrESS phenotype (Figure 2B). However, cells treated during mitosis displayed the PrESS phenotype, with CRIB3x-GFP present at the cell ends and absent at the site of failed cell separation (Figure 2B, arrows). 3 hours after LatA washout, cells which were treated in G2 or G1/S had completed cell separation and had entered the next cell cycle. The cells treated in mitosis also entered the next cell cycle, however the original division site still failed to separate and did not display CRIB-3xGFP (Figure 2B, asterisk). These findings suggest that the PrESS phenotype is cell-cycle-dependent and only occurs in cells that are in mitosis at the time of LatA treatment. Thus, in an asynchronous population, only mitotic cells display the PrESS phenotype upon recovery from LatA treatment.

**Figure 2.**
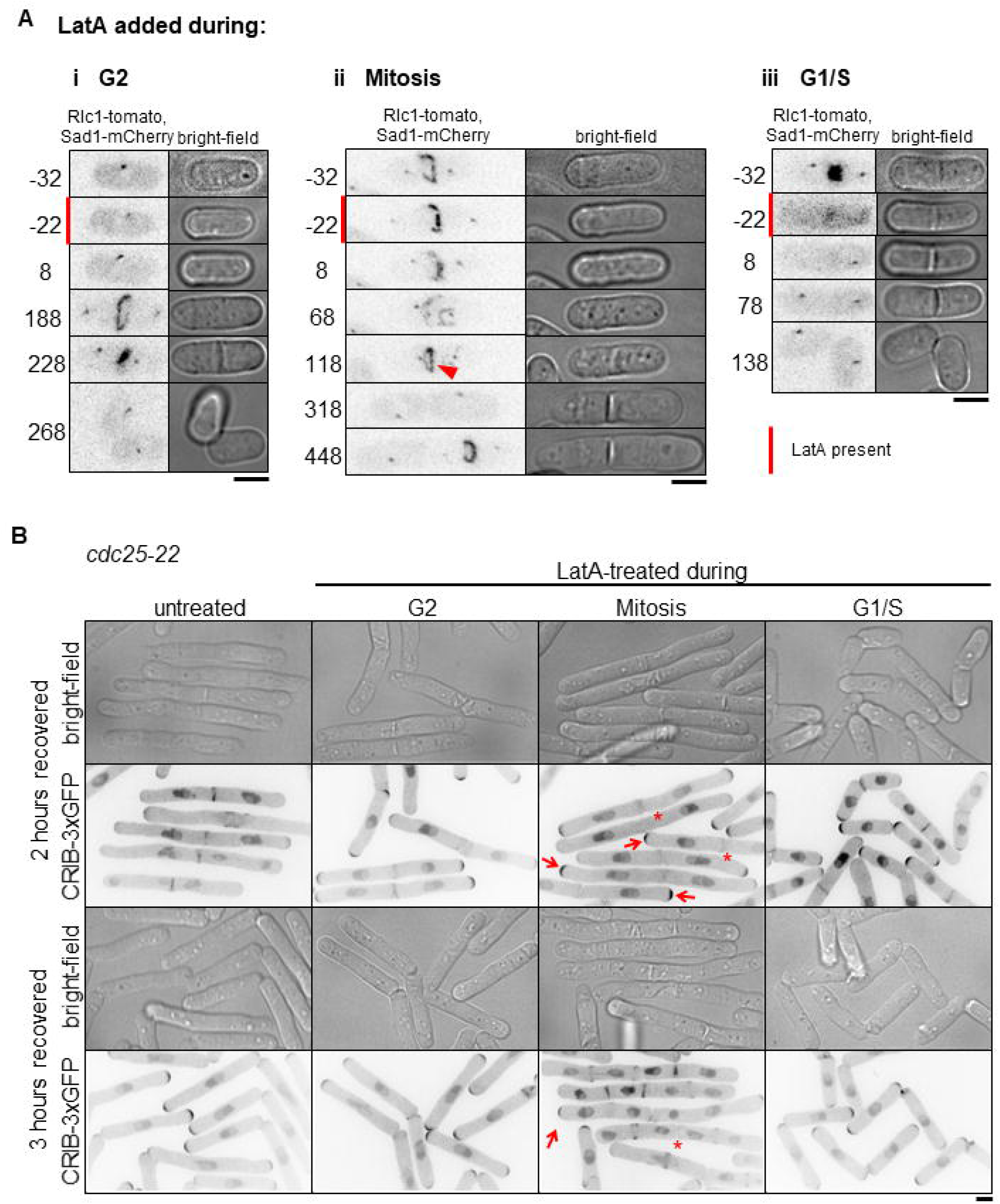
The PrESS phenotype is cell-cycle-dependent. **A)** Time-lapse imaging of cells undergoing LatA treatment and recovery. Vertical red bars denote cells undergoing a 30min 10µM LatA treatment. Time 0 marks LatA washout. Rlc1-tdTomato and Sad1-mCherry mark the actomyosin ring and spindle pole bodies and indicate the cell cycle stage of the cells. Cells in G2 (i) or in G1/S (iii) at the time of LatA treatment do not show the PrESS phenotype during recovery. Cells in mitosis at the time of LatA treatment (ii) show the PrESS phenotype during recovery. The arrowhead marks the recovered actomyosin ring undergoing constriction. **B)** *cdc25- 22* cells containing CRIB-3xGFP were synchronized by cell cycle block and release. During the indicated cell cycle stages, occurring 0 (G2), 40 (Mitosis), and 80 (G1/S) minutes after release, cells were treated for 30 minutes with LatA and then washed. Cells recovering from LatA treatment in G2 or G1/S do not yield PrESS cells while those undergoing mitosis exclusively yield PrESS cells. Arrows mark PrESS cells with active Cdc42 at cell ends. Scale bars, 5μm.

### PrESS cells activate Cdc42 at the growing ends at the expense of the division site

Why do PrESS cells fail to separate? Cell separation requires the formation of a tri-layered septum which is subsequently digested by glucanases (Cortes et al., 2016; Garcia Cortes et al., 2016; Sipiczki, 2007). The septum is comprised of a primary septum flanked by secondary septum. Glucanases are delivered to the cortex at the outer edge of the septum barrier to digest the primary septum and allow separation. Separation defects occur either due to inadequate primary septum formation or improper delivery of the glucanases. We asked if separation failure in PrESS cells is due to improper primary septum formation. To test this, we performed electron microscopy of cells recovering from LatA treatment. Electron microscopy shows that the non-separating septum in a PrESS cell is constructed properly, displaying a distinct tri-layer similar to that observed in untreated cells (Supplemental Figure S3A). These cells also showed proper recruitment of the primary septum-synthesizing enzyme Bgs1 during septation (Supplemental Figure S3B). Moreover, the original septum in a PrESS cell often goes on to separate, albeit after a prolonged delay (Supplemental Figure S3C). The septum therefore appears competent for separation, suggesting that cell separation does not fail in PrESS cells due to a structural defect in the septum.

Cell separation failure can occur due to improper delivery of the septum-digesting glucanases to the division site (Garcia Cortes et al., 2016). Normally, once delivered to the division site, glucanases are secreted to the outer edge of the membrane barrier, where they digest the primary septum. We asked if PrESS cells’ failure to undergo cell separation is due to improper delivery of these glucanases. To test this, we imaged the glucanases Eng1 and Agn1 (Dekker et al., 2004; Martin-Cuadrado et al., 2003) at the division site of PrESS cells. In untreated cells, Eng1 and Agn1 localize to the outer edge of the membrane barrier and thus appear as a ring when viewed laterally (Supplemental Figure S3D) (Martin-Cuadrado et al., 2005). In PrESS cells, Eng1 and Agn1 appear as a patchy disc all over the membrane barrier and do not localize to the outer edge (Supplemental Figure S3D). This altered localization pattern is similar to that observed in mutants which fail to deliver glucanases and thus fail to undergo cell separation (Martin-Cuadrado et al., 2005; Perez et al., 2015; Santos et al., 2005; Wang et al., 2015). This suggests that cell separation failure in PrESS cells is due to improper delivery of the glucanases required for septum digestion.

We have recently reported that the glucanases do not localize properly to the outer edge of the membrane barrier in a hypomorphic mutant of the small GTPase Cdc42 (Onwubiko et al., 2020). Cdc42 activity stops at the cell ends during mitosis and transitions to the division site during cytokinesis (Merla and Johnson, 2000; Rincon et al., 2007; Wei et al., 2016). Once the cell separates, Cdc42 activity returns to the cell ends and growth resumes (Figure 3A, upper panel, asterisks). Competition for Cdc42 activity occurs between the two growing ends of a fission yeast cell (Das et al., 2012). Normally, Cdc42 is not activated at the old ends of daughter cells until after cell separation has occurred (Figure 3A). However, in PrESS cells, we find that Cdc42 activation occurs earlier at these ends (Figure 3B, asterisks), while it gradually fades away from the intact septum (Figure 3B, arrowhead). We quantified active Cdc42, as determined by CRIB-3xGFP intensity at the cell ends, normalized to the cell middle (Figure 3C). In non-PrESS cells, Cdc42 activity was measured at the cell middle and the ends throughout cytokinesis through cell separation. We find that Cdc42 activity is higher at the cell middle than at the cell ends throughout cytokinesis. However, in PrESS cells, Cdc42 activation at the cell ends increases over time, soon overcoming the cell middle. We hypothesize that this is indicative of a competition in PrESS cells between the ends and the septum. These sites ordinarily do not attempt to activate Cdc42 concurrently, and thus do not compete with each other. However, since PrESS cells experience a cytokinetic delay, the ends and the septum attempt to activate Cdc42 simultaneously, resulting in a competition for Cdc42 activity. The site of cell division fails in this competition, resulting in cell separation failure. These conjoined cells eventually enter the next cell cycle and show normal Cdc42 activation at the cell ends during interphase and at the new division site (arrow) during cytokinesis, as evident in the two sister cells at 120 minutes in Figure 3B showing that subsequent cell cycles progress normally.

**Figure 3.**
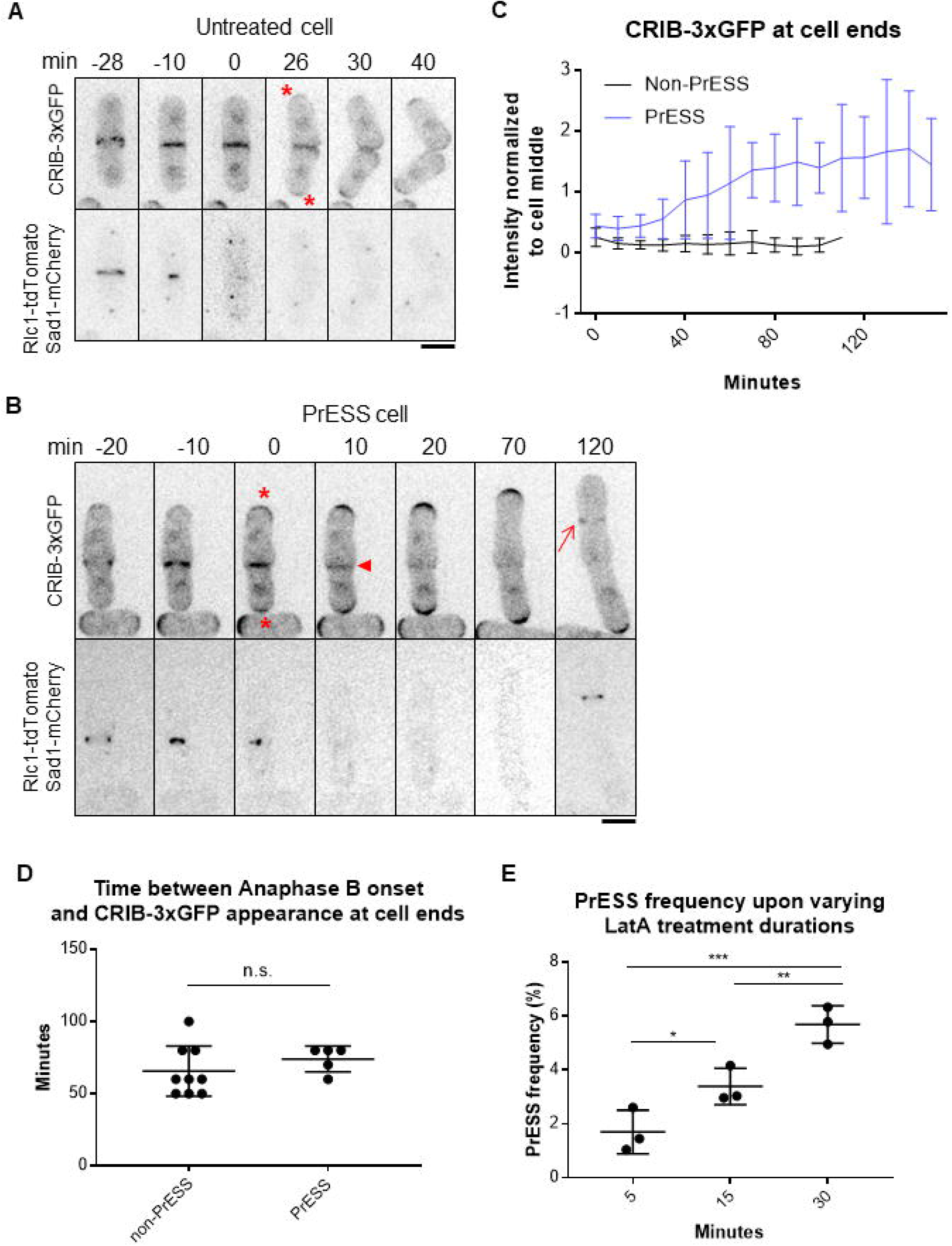
Cdc42 is activated at the ends prior to cell separation in PrESS cells. Localization of the active Cdc42 marker CRIB-3xGFP in an untreated cell (**A**) and a PrESS cell (**B**). Asterisks denote growth onset. Arrowhead denotes loss of CRIB-3xGFP from the cell middle. Arrow shows the formation of a new ring in the next cell cycle. Rlc1-tdTomato and Sad1-mCherry mark the actomyosin ring and spindle pole bodies and indicate the cell cycle stage of the cells. Time 0 marks ring/septum closure. A single z-plane is shown for clarity and hence the spindle pole body is not visible in all the time frames. **C)** Quantification of CRIB-3xGFP intensities at cell ends of non-PrESS and PrESS cells, normalized to the cell middle. n = 6 cells. **D)** The time between completion of Anaphase B and appearance of CRIB-3xGFP at the cell ends is similar in non-PrESS and PrESS cells. **E)** The PrESS frequency is lower in cells treated with LatA for a shorter duration as compared to those treated for a longer duration. All cells were treated with 10µM LatA. n > 800 cells.

We observe that in PrESS cells, Cdc42 is activated at the cell ends prematurely with respect to cytokinesis. This raises the question of whether an intrinsic timer, independent of cytokinesis, determines when Cdc42 is activated at the cell ends after division. To address this, we measured the timing of when CRIB-3xGFP returns to the cells ends after completion of anaphase B. In non-PrESS cells CRIB-3xGFP appears at the ends about 66 minutes after anaphase B. In PrESS cells, after accounting for the 30-minute LatA treatment and 60 minutes recovery, we find that CRIB-3xGFP appears at the ends about the same time (74 minutes, p = 0.3364) as non-PrESS cells (Figure 3D). This indicates that an intrinsic timer within the cell determines when Cdc42 activity should return to the cell ends after division. Cells that are undergoing this triggering event when treated with LatA will eventually show the PrESS phenotype. We thus speculate that fewer cells will undergo this Cdc42 activation trigger upon a shorter LatA treatment, resulting in fewer PrESS cells. Indeed, we find that the frequency of the PrESS phenotype does decrease with decreasing duration of LatA treatment (Figure 3E).

If Cdc42 activation indeed resumes at the cell ends at the expense of the division site in PrESS cells, we should see this reflected in the levels of Cdc42 regulators in these cells. We analyzed the localization of the Cdc42 GEF Scd1 and its scaffold Scd2 (Chang et al., 1994). During cytokinesis, Scd1 and Scd2 localize to the division membrane barrier and remain there until cell separation (Hercyk et al., 2019b; Hirota et al., 2003). In PrESS cells, we find that Scd1-3xGFP and Scd2-GFP are lost from the division site (Figure 4A, i, ii arrowheads) and appear at the cell ends, where they become enriched over time (Figure 4A, i, ii, asterisks). While it is not easy to image Scd1-3xGFP over long periods of time due to bleaching, we were able to perform time- lapse imaging of Scd2-GFP. Quantification of Scd2-GFP over time reveals a localization pattern similar to that of CRIB-3xGFP, with PrESS cells showing an increase in signal intensity at the cell ends in relation to the cell middle (Figure 4Bi).

**Figure 4.**
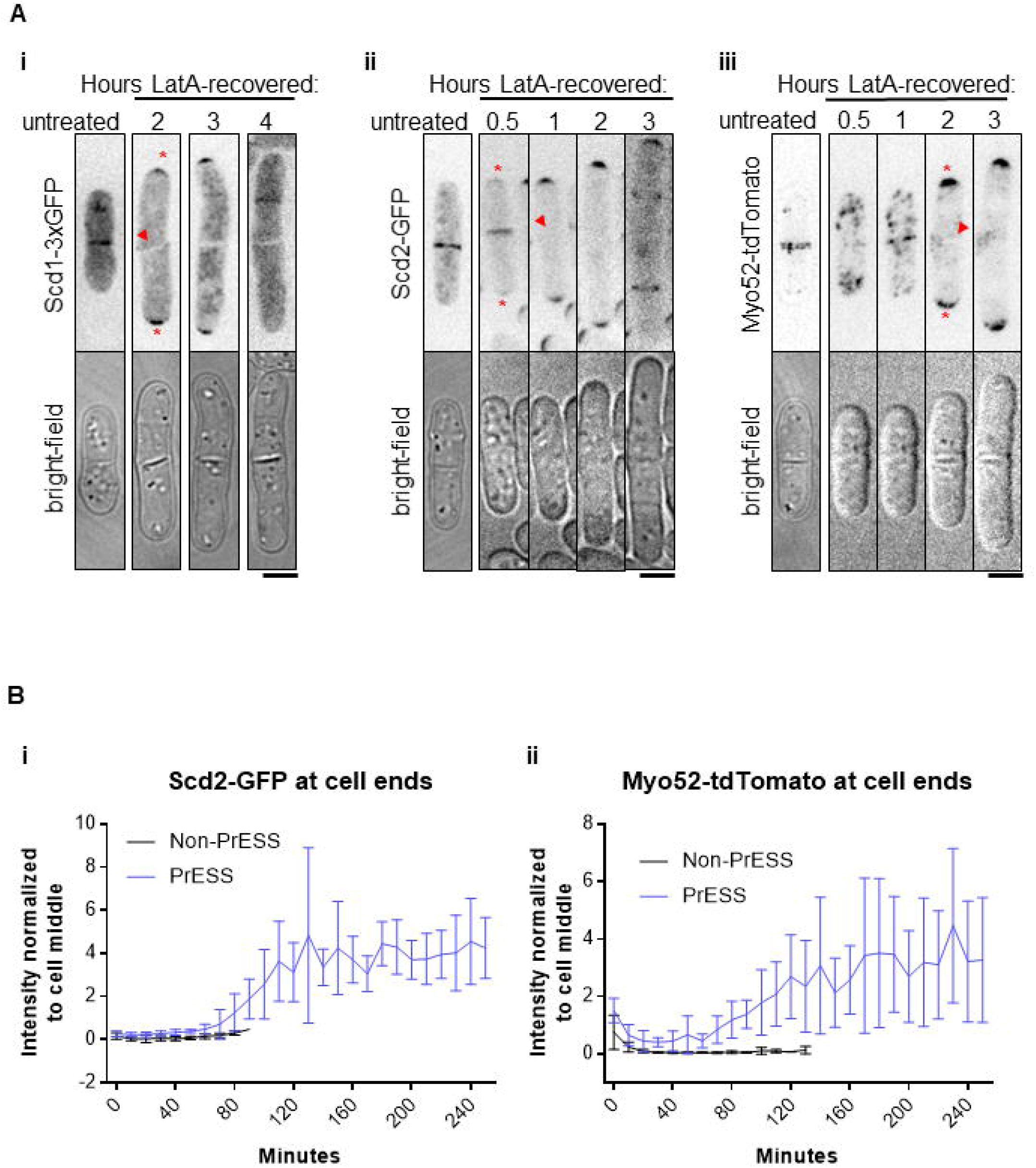
Cdc42 effectors localize to the growing ends of PrESS cells. **A)** In untreated cells, Scd1-3xGFP (i), Scd2-GFP (ii), and Myo52-tdTomato (iii) localize to the septum. When PrESS cells start to grow, all three gradually disappear from the cell middle (arrowheads) and appear at the growing cell ends (asterisks). Maximum projections of Myo52-tdTomato and single planes of Scd1-3xGFP and Scd2-GFP are shown. For Scd1-3xGFP images were taken every hour after LatA washout and different cells at the depicted time point are shown. Time lapse imaging after LatA washout was performed for Scd2-GFP and Myo52-tdTomato. A different untreated Scd2-GFP or Myo52-tdTomato is shown while treated cells shown are the same cell over time **B)** Quantification of Scd2-GFP (i) and Myo52-tdTomato (ii) intensities at cell ends of non- PrESS and PrESS cells, normalized to the cell middle. n = 6 cells. Scale bars, 5μm.

Our data show that in PrESS cells, glucanases required for cell separation are not trafficked properly during cytokinesis. Membrane trafficking occurs at sites of Cdc42 activation (Estravis et al., 2012; Estravis et al., 2011; Harris and Tepass, 2010; Murray and Johnson, 2001). Thus, in PrESS cells, with Cdc42 activity decreasing at the cell middle and increasing at the ends, we expect a similar pattern for the membrane trafficking apparatus. To test this, we analyzed the recruitment of the type 5 myosin Myo52 to the division site (Win et al., 2001). Indeed, in PrESS cells, we observe that Myo52-tdTomato intensity gradually increases at the cell ends while it decreases at the septum (Figure 4Aiii, Bii). Myo52-tdTomato levels at the cell ends over time thus resemble that of CRIB-3xGFP and Scd2-GFP in PrESS cells (Figure 3C, 4B). These findings together suggest that, in PrESS cells, Cdc42 activity, and consequently the membrane trafficking apparatus, transition to the cell ends from the division site. This leads to cell separation failure due to improper delivery of the glucanases that are supposed to digest the primary septum.

### A candidate screen to identify cell-cycle-dependent regulators of Cdc42 activation

Next, we wanted to identify how Cdc42 is regulated in a cell-cycle-dependent manner at the cell ends. We hypothesized that a Cdc42 regulator is under cell cycle control. Once cell division completes, this regulator allows Cdc42 activation at the ends. To identify this regulator, we performed a candidate screen of known mutants of Cdc42 regulators, listed in Table S2. We measured the frequency of the PrESS phenotype in these mutants. We considered the different mutants’ cell sizes in computing the PrESS frequency, as described in the Materials and Methods. We observed the PrESS phenotype in all but one of the mutants analyzed. The PrESS phenotype did not occur in the *cdc16-116* mutant (Table S2). Cdc16 is a GAP that inactivates the SIN pathway. This hypomorphic *cdc16-116* mutant shows constitutive SIN activation that leads to repeated septation and absence of cell growth (Schmidt et al., 1997). This suggests that the PrESS phenotype occurs as long as a cell is capable of growth. It is reported that Cdc42 activation in LatA-treated cells is regulated by the MAP kinase Sty1 (Mutavchiev et al., 2016; Toda et al., 1996). We asked if the PrESS phenotype is a Sty1-dependent stress response. The PrESS phenotype persisted in *sty1Δ* mutants, suggesting that this phenotype is not a stress response to LatA treatment (Supplemental Figure S4A). Cell polarity in fission yeast depends on actin as well as the microtubule cytoskeleton. The microtubule-dependent Tea1 protein localizes polarity markers to the cell ends (Feierbach et al., 2004; Glynn et al., 2001). However, *tea1Δ* mutants show the PrESS phenotype after recovery from LatA treatment. This suggests that the PrESS phenotype is independent of the microtubule-associated polarity markers (Supplemental Figure S4B).

Polarized Cdc42 activation depends on Ras1 GTPase, which promotes polarized localization of the Cdc42 GEF Scd1 (Chang et al., 1994; Chen et al., 2019; Lamas et al., 2020c). However, *ras1Δ* mutants continue to display the PrESS phenotype after recovery from LatA treatment (Supplemental Figure S4C). We also find that loss of either of the partially redundant GEFs, Scd1 or Gef1, does not lead to reduced PrESS frequency (Figure 5A, B). This suggests that cell-cycle-dependent activation of Cdc42 at the ends is not dependent on a specific GEF, and that as long as a GEF is available, Cdc42 will be activated at the cell ends. We find that *gef1Δ* shows a higher PrESS frequency than wild type (Figure 5B). We have previously shown that in *gef1Δ* cells, the old end competes with the new end for Cdc42 activity more effectively than wild type (Hercyk et al., 2019b). Thus, in *gef1Δ* mutants, a robust old end can explain the higher PrESS frequency we observe.

**Figure 5.**
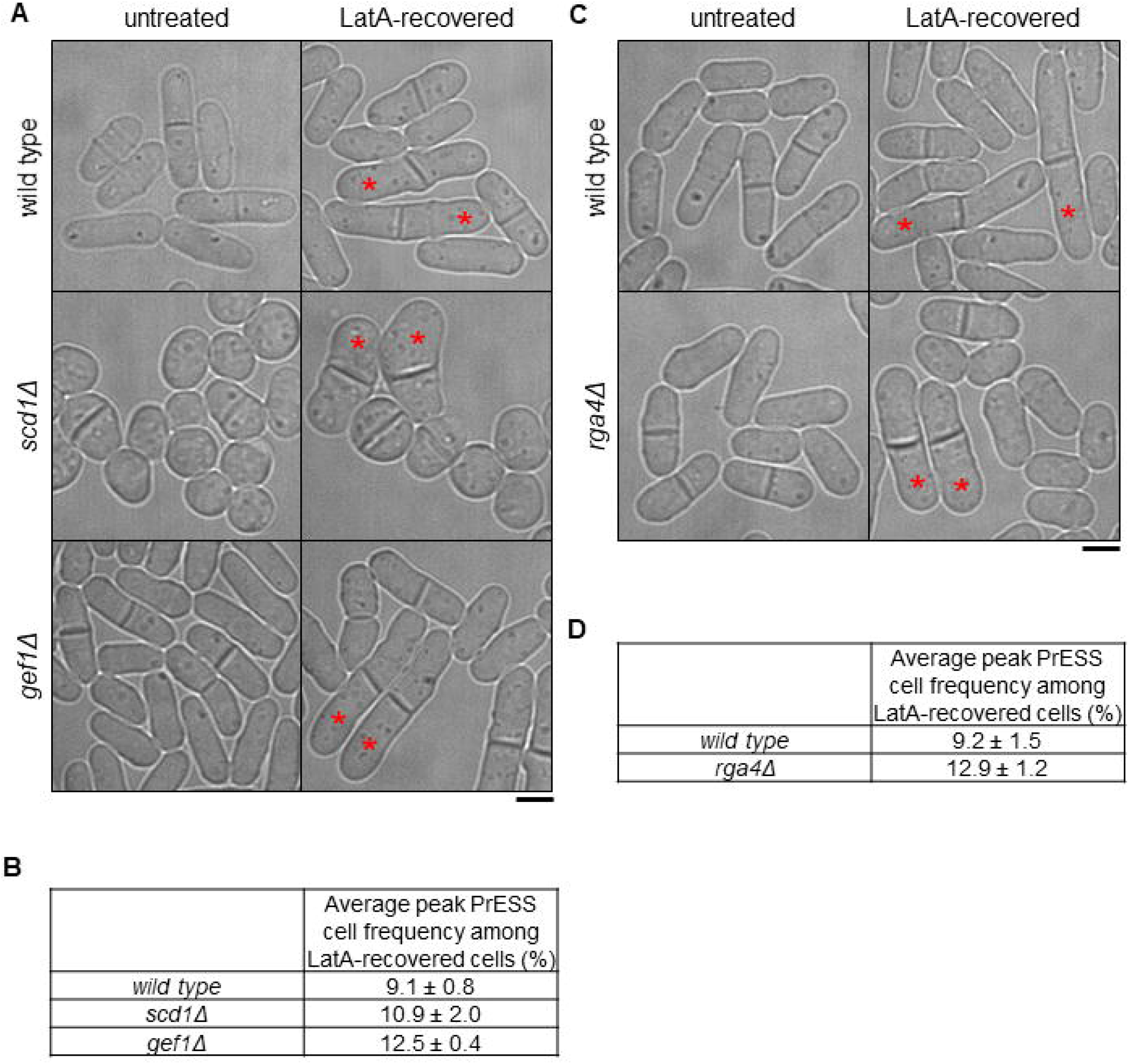
Loss of negative regulation of Cdc42 increases the incidence of the PrESS phenotype. **A, B)** PrESS phenotype persists in mutants of Cdc42 activators *scd1Δ* and *gef1Δ*. *scd1Δ* does not show a significant change from wild type, while *gef1Δ* shows an increase (p = 0.015) (Ordinary 1-way ANOVA with Dunnett’s multiple comparisons test). Asterisks denote PrESS cells. Repeated in triplicate, n ≥ 350 cells per experiment. **C, D)** Loss of Cdc42 inactivator Rga4 increases PrESS frequency during recovery from LatA treatment (Student’s t-test, p = 0.033). Repeated in triplicate, n ≥ 350 cells per experiment. Scale bars, 5μm.

Next, we analyzed the PrESS phenotype in deletion mutants of the Cdc42 GAPs Rga4 and Rga6. These GAPs inactivate Cdc42 at the cortex (Das et al., 2007; Revilla-Guarinos et al., 2016; Tatebe et al., 2008). We did not see any change in the PrESS frequency of *rga6Δ* cells. In contrast, *rga4Δ* shows a significantly higher incidence of the PrESS phenotype compared to wild type cells (p < 0.05) (Figure 5C, D). *rga4Δ* cells are wider and shorter than wild type (Das et al., 2007). This observation is consistent among PrESS cells. One explanation for the higher PrESS frequency in *gef1Δ* and *rga4Δ* cells could be a longer mitotic phase or septation defect. However, mitosis is not prolonged, compared to wild type cells, in either *gef1Δ* mutants (Wei et al., 2016) or in *rga4Δ* mutants (Supplemental Figure S5C). We did observe a higher septation index in *gef1Δ* mutants but not in *rga4Δ* mutants (Supplemental Figure S5A, B). This indicates that the higher PrESS frequency in these mutants is not due to a mitotic delay or a septation defect.

### Rga4 localization is regulated in a cell-cycle-dependent manner

In PrESS cells, Cdc42 is activated at the ends earlier with respect to septation. This enhanced Cdc42 activity at the ends could be due to either an increased activity of one of its GEFs or loss of one of its GAPs. We find that the GEFs are non-essential for the PrESS phenotype, suggesting that increased GEF activity is not responsible for this phenotype. Since deletion of the Cdc42 GAP *rga4* results in a higher PrESS frequency, we asked if Rga4 has a role in regulating Cdc42 activity at the ends and thus growth during mitosis. Previous studies have shown that Rga4 localizes mainly to the cell sides, where it blocks Cdc42 activation (Das et al., 2007; Tatebe et al., 2008). In wild type cells, we observe that Rga4-GFP localizes mostly to the cell sides during interphase (Figure 6A). As the cell enters division, Rga4-GFP can be found not only at the cell sides, but also at the ends (Figure 6A, asterisks). Time-lapse imaging of Rga4- GFP (10 sec intervals for 1 minute) indicates that Rga4-GFP at the cell ends is dynamic in nature (Supplemental Figure S6A). A time-lapse projection of Rga4-GFP (10 sec intervals for 5 minutes) shows increased signal at the cell ends in mitotic cells as compared to interphase cells (Figure 6B, asterisks; Supplemental Movie S3). We measured the mean intensity of Rga4-GFP at cell ends in these time-projected wild type cells. We find that mitotic cells have significantly more Rga4-GFP at their cell ends than cells in G2 or G1/S (Figure 6C). We also observed that in time-projected cells, the intensity of Rga4-GFP at the cortex in relation to the cytoplasm is elevated in G2 and G1/S cells compared to mitotic cells (Supplemental Figure S6B).

**Figure 6.**
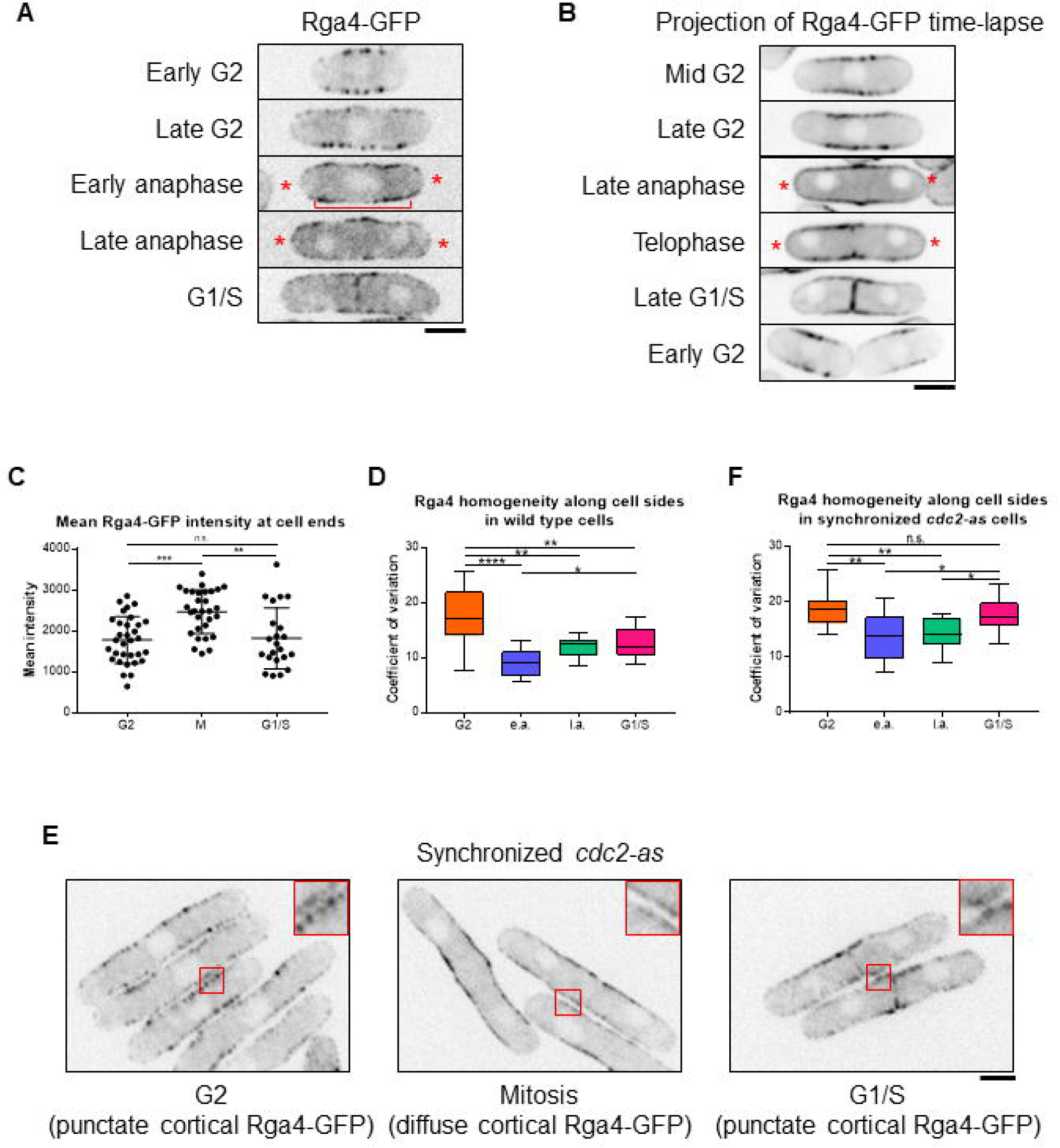
Rga4 localization changes in a cell-cycle-dependent manner. **A)** During G2 and septation (G1/S), Rga4-GFP localizes to the cortex along the cell sides as distinct puncta. During mitosis (late and early anaphase) Rga4-GFP at the cortex appears diffuse (bracket) with localization extending to the cell ends (asterisks). **B)** Time-lapse projection (10 sec intervals over 5 mins) of Rga4-GFP localization at different cell cycle stages. Asterisk denotes enhanced tip localization in mitotic cells. **C)** Quantification of Rga4-GFP intensities to cell ends during different cell cycle stages. Significantly more Rga4-GFP localizes to cell ends during mitosis than during either G2 (p = 0.0002) or G1/S (p = 0.0027). **D)** Quantification of the Rga4-GFP localization pattern along the cell sides (increased co-efficient of variance = decreased homogeneity = increased punctate appearance) in wild type cells through different cell cycle stages. (n ≥ 10 cells) e.a. = early anaphase; l.a. = late anaphase. **E)** *cdc2-as* mutant cells blocked in G2 upon 1NM-PP1 treatment show distinct Rga4-GFP puncta along the cell sides. After 1NM-PP1 washout, *cdc2-as* mutant enters mitosis and Rga4-GFP appears more diffuse along the cortex. Once in G1/S, Rga4-GFP again localizes as puncta. **F)** Quantification of the Rga4-GFP homogeneity along the sides, as done for D, in synchronized *cdc2-as* cells through different cell cycle stages. (n ≥ 14 cells) Ordinary one-way ANOVA with Tukey’s multiple comparisons was used for statistical analysis. Scale bars, 5µm.

We further observe that not only the localization, but also the distribution, of Rga4-GFP along the cortex changes throughout the cell cycle. In G2, Rga4-GFP appears as puncta dispersed along the cell sides (Figure 6A). During mitosis, it appears to be spread more homogeneously along the cortex (Figure 6A, bracket). We quantified these changes by measuring the coefficient of variation of Rga4-GFP distribution along the cortex. The higher the coefficient of variation, the less homogeneous the distribution of Rga4-GFP. We find that the coefficient of variation is higher during G2 than it is during mitosis, reflecting our observation that Rga4-GFP appears more punctate during G2 and more diffuse during mitosis (Figure 6D). The coefficient of variation drops in cells undergoing mitosis and appears to begin to increase again in G1/S cells. This suggests that, in mitotic cells, the distribution of Rga4-GFP along the cortex is more homogeneous, and gradually reverts back to a more punctate appearance as the cells enter G1/S.

Our analysis indicates that Rga4 localization and distribution change through different cell cycle stages. We asked if these changes are indeed cell-cycle-dependent. To test this, we analyzed the localization pattern of Rga4-GFP throughout the cell cycle via cell-cycle block and release. In our experience Rga4-GFP localization shows artifacts at high temperature and hence we did not use the *cdc25-22* mutants. Instead we used an analog-sensitive mutant *cdc2-as* of the mitotic kinase Cdk1 which allows specific inhibition of its kinase activity upon addition of the inhibitor 1NM-PP1 (Aoi et al., 2014). In untreated and thus asynchronous *cdc2-as* cells, the Rga4-GFP distribution pattern is similar to that observed in control cells (Supplemental Figure S6C). Next, we synchronized *cdc2-as* cells by adding the inhibitor 1NM-PP1. We distinguished between different cell cycle stages by observing the nucleus/nuclei and Rga4 localization at the division site (Supplemental Figure S6D). After arresting the cells in G2, we washed out the inhibitor to allow release into mitosis. Distribution of Rga4-GFP along the cortex was analyzed in cells at different cell cycle stages. Cells arrested in G2, when Cdc2 activity is inhibited, show a high coefficient of variation, corresponding to a more punctate Rga4-GFP distribution (Figure 6E, F). Approximately 20 minutes after removal of the inhibitor, the cells were in early anaphase. Under these conditions, the coefficient of variation of Rga4-GFP distribution is significantly reduced, corresponding to a more homogeneous distribution (Figure 6E, F). Similarly, cells in late anaphase, about 40 minutes after release, also show a decreased coefficient of variation of Rga4-GFP distribution. The coefficient of variation in *cdc2-as* cells synchronized in G1/S (∼70 minutes after release) is higher than in mitotic cells, suggesting that the punctate distribution reappears after mitosis. However, in asynchronous cells, the coefficient of variation in G1/S is lower than it is in G2 (Supplemental Figure S6C). During G1/S, Rga4-GFP distribution returns to a more punctate appearance, similar to G2. This change is likely better captured in synchronized cells. In cell-cycle-arrested *cdc2-as* cells, which are larger than wild type cells, Rga4-GFP was not clearly visible at the cell ends. It is possible that the increased cell size of these arrested cells leads to a decrease in the local concentration of Rga4-GFP at the ends.

### Loss of Rga4 allows polarized growth activation after cell division

Our data suggest that Cdc42 regulation at the cell ends is cell-cycle-dependent and that loss of *rga4* enhances PrESS frequency. Moreover, we find that localization of Rga4 to cell ends increases during mitosis when Cdc42 activation at those ends is known to decline. We propose that two elements are necessary for polarized growth to occur: MOR pathway activation leading to protein synthesis, as well as Cdc42 activation at the cell ends. During mitosis, the MOR pathway is inactive and Rga4 is present at the ends, and growth does not occur. As cell division completes, the MOR pathway becomes active and the cell ends lose Rga4, allowing local Cdc42 activation. In PrESS cells, cytokinesis is delayed, but the MOR pathway and Cdc42 at the ends are activated as normal, resulting in polarized growth. We should thus be able to recapitulate the PrESS phenotype in cells both constitutively activating the MOR pathway and activating Cdc42 at the ends even without a delay cytokinesis. Constitutive activation of the MOR pathway via expression of the *nak1-mor2* fusion leads to premature protein synthesis during cytokinesis (Gupta et al., 2014). This includes the glucanases that are then prematurely delivered to the incomplete septum barrier, resulting in cell lysis (Figure 7A, arrows; Supplemental Figure 7A). We hypothesized that if the cell ends were instead able to activate growth, the glucanases would be delivered to the cell ends rather than the division site, resulting in a PrESS-like phenotype. Based on our previous observations, we should accomplish this by deleting *rga4* in a mutant constitutively activating the MOR pathway. Since *nak1-mor2*-expressing cells undergo cell lysis resulting in cell death, they display a low optical density (Gupta et al., 2014). We observed cell lysis in *rga4^+^* cells with *nmt1-nak1-mor2* under both expressing and repressing conditions. This is likely due to leaky expression of *nak1-mor2* in these cells. However, cell lysis was enhanced under expressing conditions compared to repressing conditions, resulting in a significantly lower optical density in *rga4^+^* cells expressing *nak1-mor2* (Figure 7B). Accordingly, the fraction of lysed cells in *rga4Δ* mutants expressing *nak1-mor2* was significantly reduced compared to *rga4^+^* cells expressing *nak1-mor2* (Figure 7C). These cells fail to undergo cell separation and continue to grow from their ends, similar to PrESS cells (Figure 7A, arrowhead). Indeed, while *rga4^+^* cells expressing *nak1-mor2* did not show PrESS-like cells, *rga4Δ* cells expressing *nak1-mor2* consistently showed this phenotype in the population (Figure 7D). Note that *rga4Δ* cells are capable of cell separation, and thus failure to separate in *nak1-mor2*-expressing conditions is due to premature growth at the ends, similar to PrESS cells.

**Figure 7.**
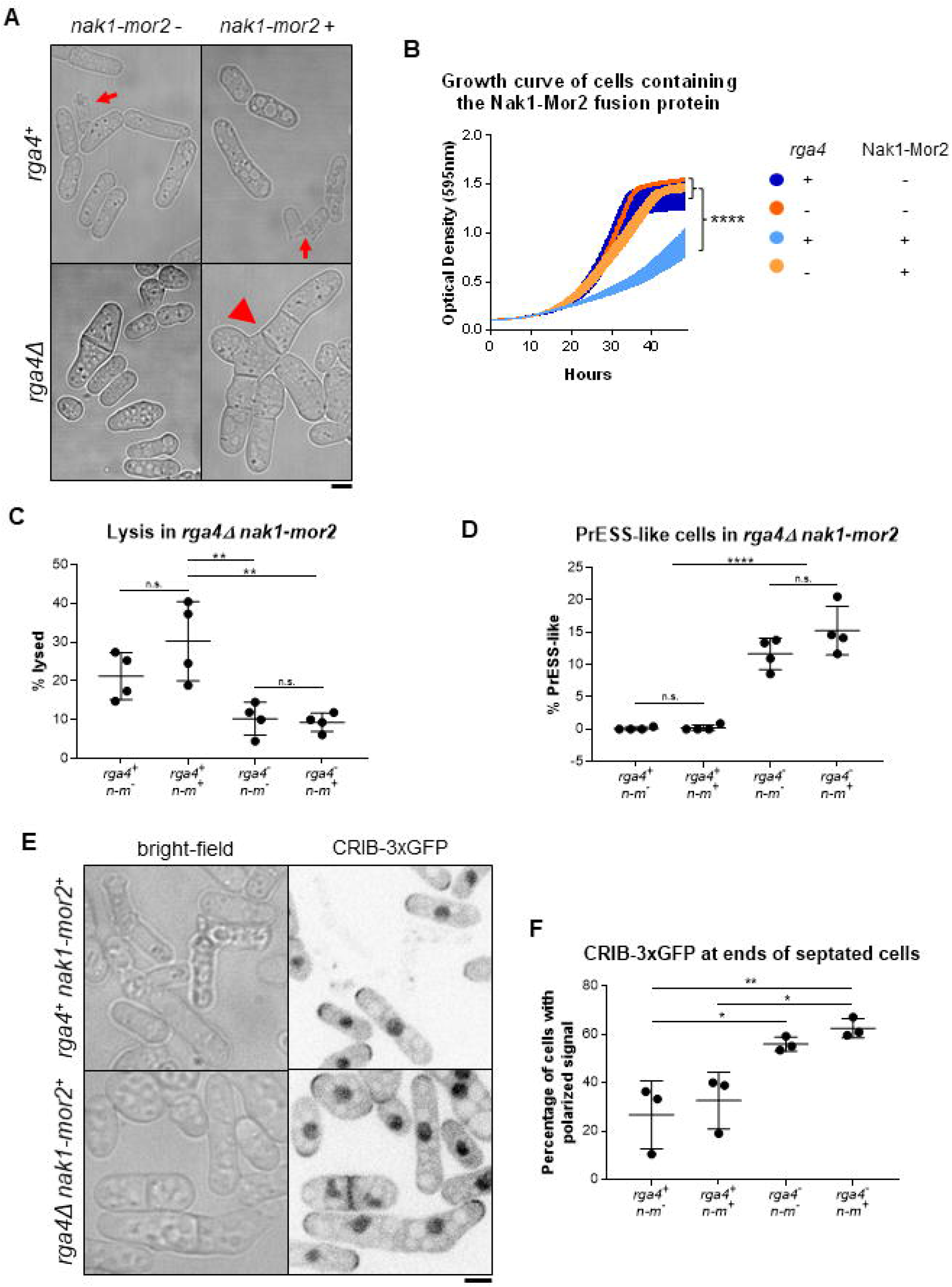
**Deletion of *rga4* allows polarized growth in cells with constitutive MOR activity. A**) Cells expressing *nak1-mor2* (-thiamine) show cell lysis (red arrow) at the division site during cytokinesis in *rga4^+^* cells, while in *rga4Δ* mutants *nak1-mor2* expression leads to a PrESS-like phenotype (arrowhead) in which cells fail separation and grow from the ends. **B)** Growth curve of *rga4^+^* or *rga4Δ* cells either expressing *nak1-mor2* or repressing *nak1-mor2.* Optical density was measured every 15 minutes at 595nm for 2 days. (Ordinary one-way ANOVA with Tukey’s multiple comparisons. **** = p < 0.0001; 5 experiments averaged for each genotype) **C)** *rga4^+^* cells expressing *nak1-mor2* (*n-m*) showed significantly more lysis than *rga4Δ* cells either repressing or expressing *nak1- mor2* (p = 0.0042; p = 0.0030) **D)** Percent of cells displaying a PrESS-like phenotype. *rga4Δ* cells containing *nak1-mor2* (*n-m*) display a significantly higher PrESS-like frequency than *rga4^+^* cells containing *nak1-mor2* (p < 0.0001). **E)** *rga4^+^* and *rga4Δ* cells expressing *nak1-mor2* and containing CRIB-3xGFP. While only a few septated *rga4^+^* cells show CRIB-3xGFP at their ends, a large fraction of septated *rga4Δ* cells do. **F)** Quantification of CRIB-3xGFP presence at cell ends in *rga4^+^* and *rga4Δ* cells expressing *nak1-mor2* (*n-m*). Ordinary one-way ANOVA with Tukey’s multiple comparisons was used for statistical analysis. Experiments repeated in triplicate. Scale bars, 5µm.

Next, we asked if this PrESS-like phenotype is due to Cdc42 activation at the ends in dividing cells. PrESS cells have a septum and show active Cdc42 at their ends. Thus, we analyzed CRIB-3xGFP at the ends of septated cells in *rga4^+^* and *rga4Δ* cells expressing *nak1-mor2*. We find that, in *rga4Δ* cells expressing *nak1-mor2*, a larger fraction of septated cells shows CRIB-3xGFP at the cells ends compared to the *rga4^+^* control (Figure 7E and F). We asked if the PrESS-like phenotype was specific to Rga4. We show that the PrESS frequency also increases in *gef1Δ* mutants (Figure 5). However, we did not observe PrESS-like cells or rescue of cell lysis in *gef1Δ* cells expressing *nak1-mor2* (Supplemental Figure S7B). Similarly, it is possible that the PrESS-like phenotype could be due to loss of any Cdc42 GAP. However, in *rga6Δ* cells expressing *nak1-mor2*, we did not observe PrESS-like cells or rescue of cell lysis (Supplemental Figure S7B). We thus propose that lack of Rga4 at the cell ends allows the ends to better compete for growth machinery, preventing localization of these proteins to the division site. This suggests that under normal conditions, after MOR pathway activation, cell growth occurs once the ends lose Rga4.

## Discussion

Although much is known about how polarized growth occurs in fission yeast, it is unclear how polarized growth resumes at the ends after cell division completes. Specifically, it is unknown how Cdc42 activity transitions from the division site to the ends to drive polarized growth. Since Cdc42 activation at the ends normally does not occur until after completion of cell separation, we asked if the timing of Cdc42 activation and consequent growth resumption could depend on cell separation itself. However, here we show that the timing of Cdc42 activation at the old end is independent of cell separation and is instead cell-cycle-dependent.

The MOR pathway promotes cell separation and polarized growth after mitosis (Chen et al., 2019; Gupta et al., 2014; Nunez et al., 2016; Ray et al., 2010). The final step in cell division, cytokinesis, involves actomyosin ring formation and subsequent constriction in coordination with septum formation and membrane invagination (Cheffings et al., 2016; Garcia Cortes et al., 2016; Pollard, 2010). After ring constriction, the septum matures to form a tri-layer – a primary septum flanked by secondary septum (Cortes et al., 2016; Garcia Cortes et al., 2016). MOR pathway activation allows synthesis of glucanases, which are delivered to the outer edge of the membrane barrier to precisely digest the primary septum, resulting in cell separation (Martin-Cuadrado et al., 2005; Nunez et al., 2016; Santos et al., 2005). Once septum digestion starts, Cdc42 activity transitions to the old ends and polarized growth initiates. It was unclear how cell separation and growth initiation occur sequentially since the MOR pathway promotes both processes. Constitutively activating the MOR pathway leads to cell lysis during division, which is caused by premature cell separation as a result of unregulated glucanase synthesis and delivery (Gupta et al., 2014). Here we report the novel PrESS phenotype, in which polarized cell growth initiates during septation. Our data suggest that, in these cells, Cdc42 is activated at the ends at the expense of cell separation. Activation of Cdc42 at the ends of these cells coincides with a decrease in Cdc42 activity at the division site.

Thus, instead of delivering glucanases to the division site resulting in premature cell lysis, the ends show polarized growth. This indicates that, under normal conditions, Cdc42 activity at the ends is inhibited while the division site successfully undergoes cell separation. The ends are able to activate Cdc42 only after cell separation completes, even though the MOR pathway is active throughout this stage. Indeed, we find that constitutively activating the MOR pathway does not lead to Cdc42 activation at the cell ends during division. These observations together suggest that, in addition to MOR activation, another pathway regulates Cdc42 at the ends after division.

Our results indicate that Rga4 is regulated at the cell ends during mitosis. We show that Rga4 localizes all the way to the cell ends during mitosis, instead of being restricted to the cell sides (Das et al., 2007). It is possible that Rga4 at the ends during mitosis blocks Cdc42 activation. Accordingly, we hypothesize that, in the absence of *rga4*, cells should initiate growth at their ends during division as long as protein synthesis via the MOR pathway is active, thus recapitulating the PrESS phenotype. Indeed, in cells constitutively activating the MOR pathway, loss of *rga4* leads to end growth and cell separation failure in a recapitulation of the PrESS phenotype, thus supporting our hypothesis.

How does Rga4 localize to the cell ends during mitosis? Rga4 typically localizes to the cell sides during G2 and is excluded from the cell ends. During this phase, Rga4 appears as distinct puncta dotted along the cell sides. However, during mitosis, Rga4 appears more diffuse along the cortex and extends all the way to the cell ends. Cells arrested in G2 show distinct punctate Rga4 distribution at the cell sides. Upon release, when the cells enter mitosis, Rga4 appears diffuse at the cortex. It is unclear how Rga4 localization and distribution are regulated during mitosis. Rga4 could be a direct target of the mitotic kinase Cdc2 or of an unknown cell-cycle-dependent regulatory module. In another study, Rga4 has been identified as a target of Cdk1 kinase activity, however the physiological significance of this interaction is unknown (Swaffer et al., 2016). We are currently investigating the nature of cell-cycle-dependent Rga4 regulation during mitosis. It is not clear how Rga4 is removed from the ends upon completion of mitosis. Rga4 still lingers at the ends after activation of the anaphase-promoting complex, which promotes Cdk1 inactivation. It is possible that Rga4 undergoes some modification that gradually leads to its loss from the ends. This could be mediated by a phosphatase, by turnover of Rga4, or by an unknown mechanism. While it is unknown how changes in the localization pattern of Rga4 affects Cdc42 regulation, a similar concept has been recently shown in another study (Gerganova et al., 2021). There, the authors show that artificially localizing a Cdc42 GAP to the cortex completely blocks Cdc42 activity, while conditions leading to its oligomerization result in its loss from the ends, enabling local Cdc42 activation.

Similar to our findings in fission yeast, another report in budding yeast has shown that the start of polarization after cell division is independent of cytokinesis (Moran et al., 2019). Moreover, it was shown in the same study that the start of polarization is earlier in mutants lacking the Cdc42 GAPs *BEM3* and *RGA2*. In yet another study, it has been shown that Cdc42 inactivation at the onset of mitosis is mediated by BEM3 (Gihana et al., 2021). Thus, it is possible that the role of GAPs in the regulation of polarization after cell division is conserved. Moreover, in budding yeast, CDKs Pho85 and Cdc28 phosphorylate the GAP Rga2 to restrain its activity and promote polarized growth (Sopko et al., 2007). It has also been shown in budding yeast that the G1-cyclin-bound CDKs promote Cdc42 activation (McCusker et al., 2007). G1-cyclin-CDK promotes Cdc42 activation via the GEF Cdc24, thus allowing bud growth. Our findings suggest a mitosis-dependent negative regulation of Cdc42 activity. It is unclear if the G1-cyclin-CDK complex also regulates Cdc42 in fission yeast.

While we find that Rga4 localizes to the cell ends during mitosis, it should be noted that loss of *rga4* alone does not allow Cdc42 activation at the ends at this stage. This is probably because, even in the *rga4* mutants, the MOR pathway is inhibited during mitosis. Cdc42 activation at the ends is not possible while the MOR pathway is inactive. Consequently, after division, Cdc42 activation at the cell ends requires both loss of Rga4 from the cell ends and MOR pathway activation. It is possible that in an *rga4Δ* mutant, since cytokinesis is not delayed, the ends do not have a sufficient advantage in activating growth because the MOR pathway is only activated during septation. In *rga4Δ* cells constitutively activating the MOR pathway, the ends do have an advantage since the MOR pathway has been active even before septation, thus leading to growth. A recent report has shown that presence of the GAPs at the cortex prevents local Cdc42 activation even upon localization of the scaffold Scd2, which recruits Scd1 to the cortex (Lamas et al., 2020b). Another preprint suggests that Cdc42 GAP levels in daughter cells determine the pattern of polarized growth in those cells (Pino et al., 2020). These reports and our findings together indicate that loss of Rga4 from the cell ends after mitosis is necessary to allow Cdc42 activation at these ends, so long as the MOR pathway is active.

The MOR pathway is necessary for cell separation as well as polarized cell growth. Once the MOR pathway is activated, cells first undergo separation and then initiate polarized growth at their ends. Based on our findings, we propose a model to describe how polarized growth is initiated after cell separation (Figure 8). Cdc42 activity at the ends declines once the cell enters mitosis and is completely lost from these sites as mitosis progresses. At this stage, Rga4 no longer appears as distinct puncta along the cell sides. Rather, it displays a diffuse distribution along the cortex which extends to the cell ends. After the metaphase checkpoint, the cell enters anaphase, and the SIN is activated while the MOR pathway is inactivated (Simanis, 2015). Polarized cell growth at the ends stops at this stage. Once mitosis is complete, the SIN is inactivated and the MOR pathway is activated, allowing synthesis of proteins necessary for cell separation and polarized growth. As the cells separate, Rga4 is gradually lost from the ends, its punctate distribution is restored, and growth initiates. In LatA-treated cells, cytokinesis is delayed while mitosis progresses. In such cells, Rga4 is removed from the cell ends on time, allowing Cdc42 activation and growth at these sites since the MOR pathway is active, thus resulting in the PrESS phenotype.

**Figure 8.**
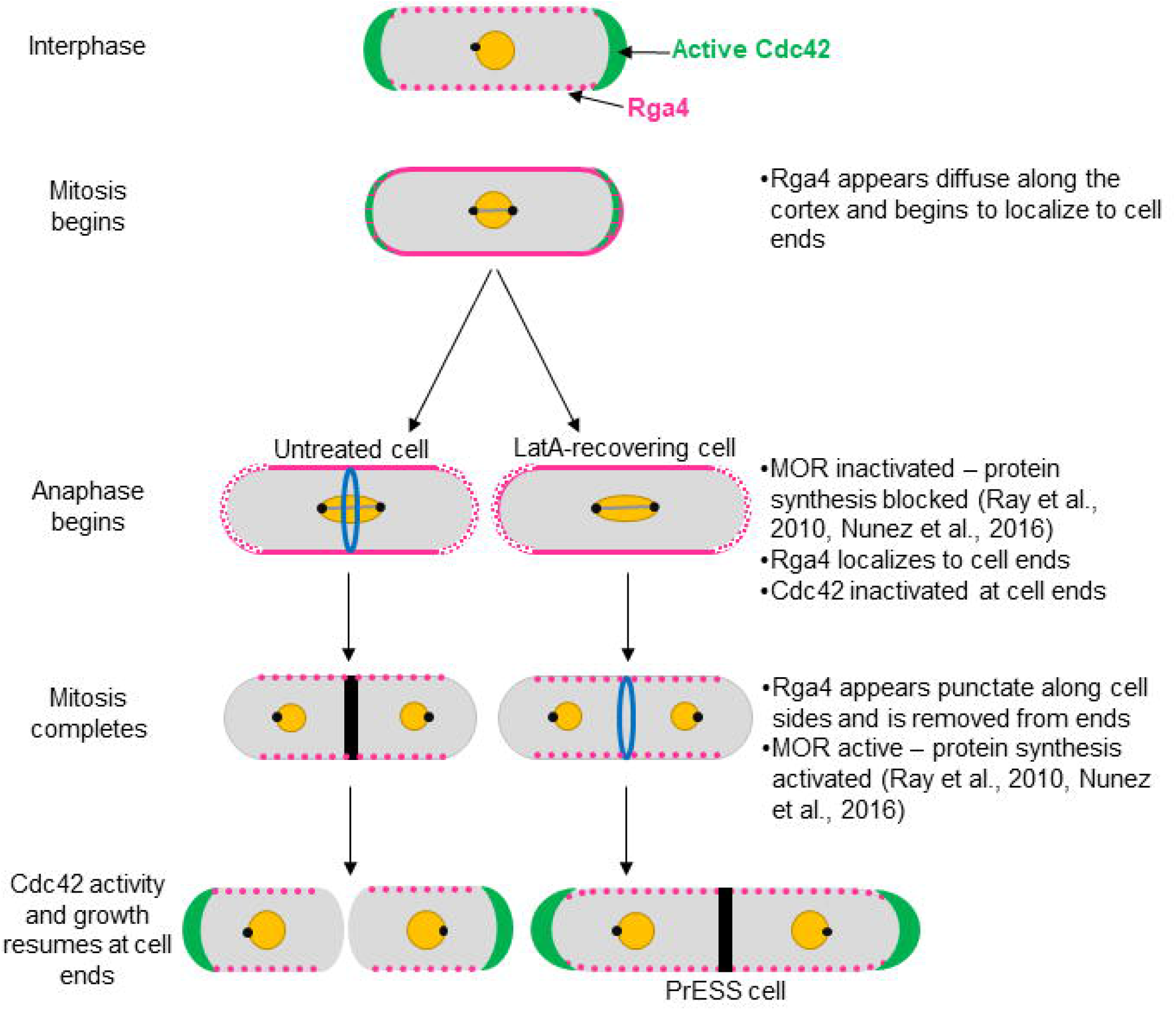
Model for growth resumption after cell division. Rga4 localizes to the cell sides during G2 (magenta). As the cells enter mitosis, Rga4 extends all the way to the cell ends (magenta). Upon activation of the anaphase-promoting complex, the MOR pathway is inactivated and Cdc42 activation and growth (green) stop at the cell ends. After mitosis, MOR is activated while Rga4 is lost from the cell ends, which now resume growth. In PrESS cells, LatA treatment delays cytokinesis while mitosis continues to proceed, and the cell progresses to interphase. In these cells, Rga4 is removed from the cell ends before the completion of cytokinesis, resulting in Cdc42 activation and growth at cell ends at the expense of cytokinetic events at the cell middle. This suggests that the timing of growth resumption upon completion of cell division is independent of cytokinesis and dependent on cell-cycle cues.

Through an artificial cytokinetic delay, we have uncovered a basic cellular principle which ensures that growth occurs at an appropriate time. It is interesting that growth is carefully suppressed during mitosis, and that a cytokinetic delay will not prevent growth. Cdc42 activity is thus regulated based on a mitotic timer without regard to completion of cytokinesis. This especially makes sense in the context of pseudohyphal growth, which fission yeast typically undergo in nature. Our findings suggest that growth initiation is cell-cycle-dependent and is timed to occur after cell separation. It is not very clear how growth and mitosis crosstalk in animal cells. A recent report suggests that, in animal cells, growth stops as cells approach the metaphase-to-anaphase transition and resumes in late cytokinesis (Miettinen et al., 2019). The Cdc42 activation pattern at cell ends in fission yeast is similar to these observations. It is possible that the nature of cell- cycle-dependent regulation of polarized growth is conserved throughout all eukaryotes. Our work provides a better understanding of how growth initiates after mitosis in fission yeast. An elegant cell-cycle-dependent system causes a subtle but effective localization change, controlling the timing of Cdc42 activation at the cell ends.

## Materials and Methods

### Strains and cell culture

The *Schizosaccharomyces pombe* strains used in this study are listed in Table S1. Cells were cultured in yeast extract (YE) medium and grown exponentially at 25°C unless otherwise specified (Moreno et al., 1991). Cells were grown for at least three days before imaging.

### Latrunculin A treatments

Cells were treated with 10µM Latrunculin A (LatA) (Sigma-Aldrich #428021) for 30 minutes. The cells were washed twice with the appropriate media, switched to a fresh tube, and washed a third time.

### PrESS frequency measurements

Wild type, *scd1Δ* and *gef1Δ* were all imaged on the same three days, and wild type and *rga4Δ* were both imaged on the same three days. Cells were LatA-treated and washed as described above, and then imaged immediately after wash as well as every hour after wash for five hours. Untreated cells were imaged as a control for each strain. The length of each septated cell was measured, and the average and standard deviation were calculated. Any cell measured to be longer than two standard deviations more than the average for the corresponding untreated sample was counted as a PrESS cell. The reported peaks are the actual PrESS percentages minus the control PrESS percentage, which was always under 1%.

### *cdc25-22* synchronization

Strains synchronized using the *cdc25-22* temperature-sensitive mutation were shifted to restrictive temperature, 36°C, for four hours and then shifted back to permissive temperature, 25°C.

### Cdc2-as inhibition and synchronization

Synchronization of cells containing *cdc2-as* was achieved via addition of 1NM-PP1 (Toronto Research Chemicals #A603003) to a concentration of 500nM for four hours. Cells were then washed three times with YE media to remove the inhibitor.

### Phalloidin staining

Phalloidin staining of actin structures was performed as described previously (Das et al., 2009; Pelham and Chang, 2002). Cells were fixed with 3.5% formaldehyde for 10 minutes at room temperature. After fixation, the cells were permeabilized with PM buffer (35 mM potassium phosphate pH 6.8, 0.5 mM MgSO4) with 1% TritonX-100, then washed with PM buffer and stained with Alexa-Fluor-phalloidin (Molecular Probes) for 30 minutes.

### Growth curve

Cells were grown at 25°C. To achieve plasmid expression, thiamine was washed out immediately before the experiment began. Cells were diluted to an O.D. of 0.01 and loaded into a 96-well plate, which was read by a BioTek Cytation 5 plate reader. O.D. measurements at 595nm were collected every fifteen minutes for 48 hours. During the experiment, the cells were maintained at 29°C, with shaking.

### Analysis of Rga4 distribution and localization

To quantify Rga4 distribution along the cell sides, a line was drawn along the cell side in ImageJ for both sides of each cell in a single plane. Then, a plot profile was generated, and the intensity values were averaged to give an average coefficient of variation for Rga4-GFP distribution along the sides.

To quantify Rga4 localization at cell ends, background subtraction was first performed on each field of cells in ImageJ. A cap was drawn around each cell end, and the intensities of the two ends of each cell were averaged together to provide one value per cell.

### Constitutive MOR activation

Cells were grown at 25°C in minimal media with thiamine. Thiamine was washed out 16 hours prior to imaging to allow plasmid expression.

### Statistical tests

GraphPad Prism was used to perform statistical tests. A Student’s t-test (two-tailed, unequal variance) was used to determine significance when comparing two samples. When comparing three or more samples, one-way ANOVA was used alongside Tukey’s multiple comparisons, unless otherwise mentioned.

### Microscopy

Imaging was performed at room temperature (23-25°C). We used an Olympus IX83 microscope equipped with a VTHawk two-dimensional array laser scanning confocal microscopy system (Visitech International, Sunderland, UK), electron-multiplying charge-coupled device digital camera (Hamamatsu, Hamamatsu City, Japan), and a 100x/1.49 numerical aperture UAPO lens (Olympus, Tokyo, Japan). We also used a spinning disk confocal microscope system with a Nikon Eclipse inverted microscope with a 100x /1.49 numerical aperture lens, a CSU-22 spinning disk system (Yokogawa Electric Corporation), and a Photometrics EM-CCD camera. Images were acquired with MetaMorph (Molecular Devices, Sunnyvale, CA) and analyzed with ImageJ (National Institutes of Health, Bethesda, MD (Schneider et al., 2012). For still and z-series imaging, the cells were mounted directly on glass slides with a #1.5 coverslip (Fisher Scientific, Waltham, MA) and imaged immediately, and with fresh slides prepared every ten minutes. Z-series images were acquired with a depth interval of 0.4 μm. For time-lapse images, cells were placed in 3.5mm glass-bottom culture dishes (MatTek, Ashland, MA) and overlaid with YE medium containing 0.6% agarose and 100 μM ascorbic acid as an antioxidant to minimize toxicity to the cell, as reported previously.

For cells attached to MatTek dishes with lectin, we first let 10μL of 1mg/1mL lectin (Sigma-Aldrich #L1395) incubate in the center of the dish at room temperature for 30 minutes. Then, we washed with 1mL of YE media three times to remove excess lectin not adhered to the dish. We spun down 1mL of cells at an O.D. of about 0.5 and removed all 950μL of the supernatant, then resuspended and pipetted the remaining 50μL into the center of the dish. We then rinsed the dish with YE media three times to remove excess cells not adhered to the lectin. YE media was subsequently kept in the dish at all times to prevent cell starvation, except during short time periods during washes and drug treatments.

### Electron microscopy

Wild type cells in late G2 were selected from a sucrose gradient. These cells were treated with 10μM LatA for 30 minutes and then washed three times with YE media. The untreated wild type control cells were not selected from a sucrose gradient. Both samples were washed in sterile water three times and fixed in 2% potassium permanganate for one hour at room temperature. They were again washed three times in sterile water and then resuspended in 70% ethanol and incubated overnight at room temperature. Dehydration was achieved via an ethanol series at room temperature: in order, two 15-minute incubations in 70% ethanol, two 15-minute incubations in 95% ethanol, and finally three 20-minute incubations in 100% ethanol. The cells were then incubated for 30 minutes in propylene oxide, followed by a 1-hour incubation in a 1:1 mixture of propylene oxide and Spurr’s resin, and lastly two 1-hour incubations in neat Spurr’s resin. The cells were pelleted via centrifugation and the supernatant carefully removed in between each incubation. The cells were transferred into small plastic tubes, and fresh Spurr’s resin was layered above to the top of the tube. The two resulting tubes were incubated at 60°C overnight. Sectioning yielded approximately 100nm sections. Sections were post-stained with lead citrate for 8 minutes. Sections were washed by dunking in three sequential beakers of sterile water thirty times each, and allowed to dry on filter paper. Samples were imaged with the Zeiss Dual Beam FIB/SEM microscope at the Joint Institute for Advanced Materials (Knoxville, Tennessee). All images were collected at 15kV.

## Acknowledgements

We thank Brian Hercyk, Udo Onwubiko, Bethany Campbell, Marcus Harrell, Samridhi Pathak, Justin McDuffie, and Abigail Vickers for discussions and feedback; Kathy Gould, Sophie Martin, Jonathan Millar, Pilar Perez, and Fulvia Verde for strains; Dannel McCollum for plasmids; and John Dunlap for electron microscopy.

This work was funded by an NIH-NIGMS R01 grant (#1R01GM136847-01) awarded to MD. JRR was awarded an NSF Graduate Research Fellowship Program (GRFP) (#1452154), as well as the NIH’s Initiative for Maximizing Student Development (IMSD) (#R25GM086761).

## Conflict of Interest

The authors declare that they have no conflict of interest.

## Notes

### Competing Interest Statement

The authors have declared no competing interest.

