## Supplemental FigureS1-S7 for "Cdc42 reactivation at growth sites is regulated by local cell-cycle-dependent loss of its GAP Rga4"

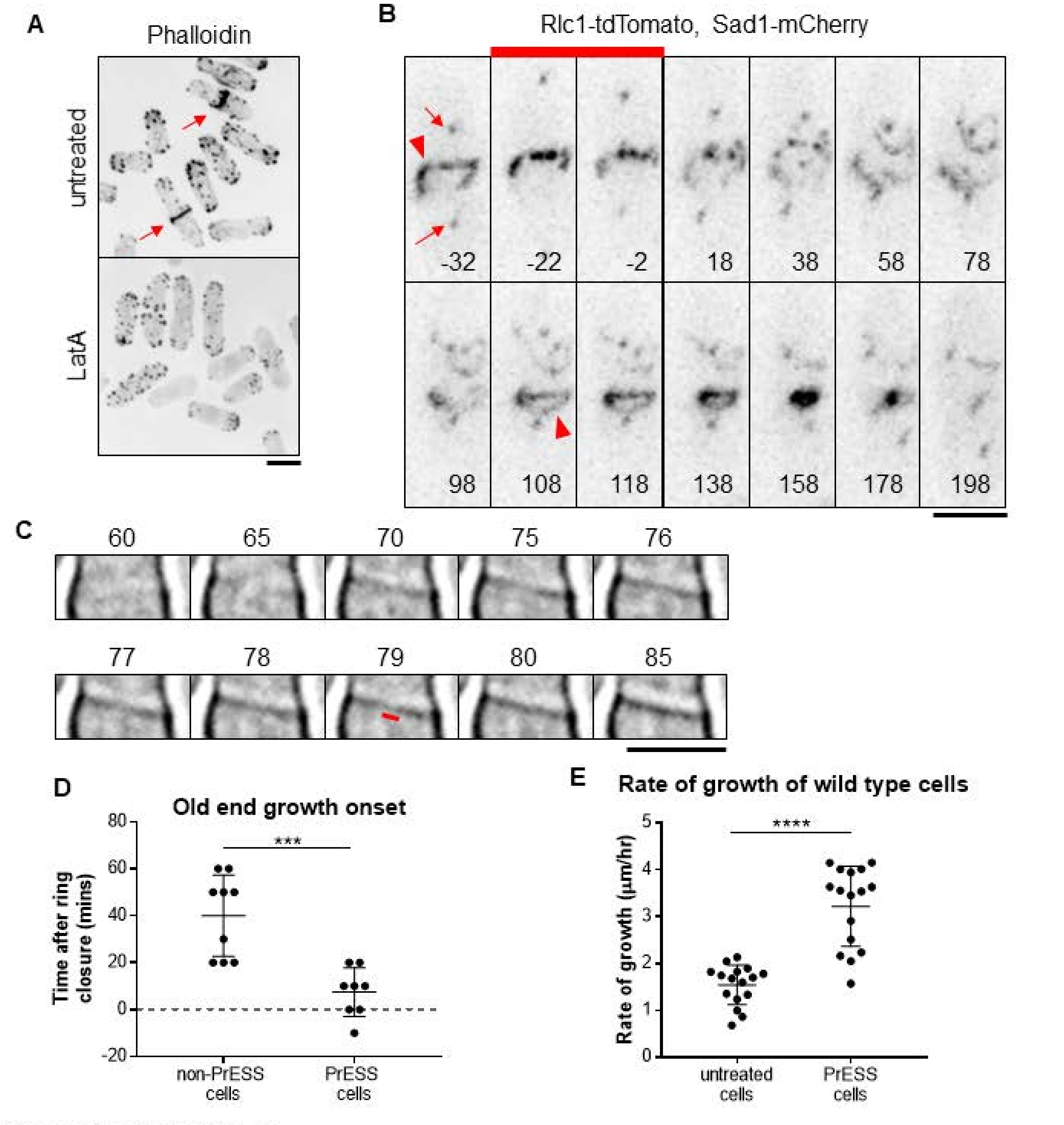


**Supplemental Figure S1**

**A)** Phalloidin staining of untreated cells and cells treated with 10µM LatA for 30 minutes. Actomyosin rings are marked by red arrows. **B)** Rlc1-tdTomato (arrowhead) and Sad1-mCherry (arrows) mark the actomyosin ring and spindle pole bodies. Time 0 is ring closure. During a 30-minute 10µM LatA treatment (**red bar**), the original actomyosin ring is disrupted. During recovery, a new ring is built, which undergoes constriction. **C)** In this sample cell, we determined that septum closure occurred at 80 minutes, because this is the first frame in which the gap that was still visible at 79 minutes (as shown by the red bar) was closed. **D)** Onset of growth in PrESS and non-PrESS cells after ring closure. **E)** Rate of old end growth of untreated and PrESS cells. ($p < 0.0001$; $n \geq 16$ cell ends. Statistical tests = Student t-test. Scale bars = 5µm.

**A**

| PrESS phenotype frequency when LatA-treated at different cell cycle stages | | | |
|---|---|---|---|
| | G2 | Mitosis | G1/S |
| % PrESS | 0 | 100 | 0 |
| n | 17 | 10 | 7 |

**B**

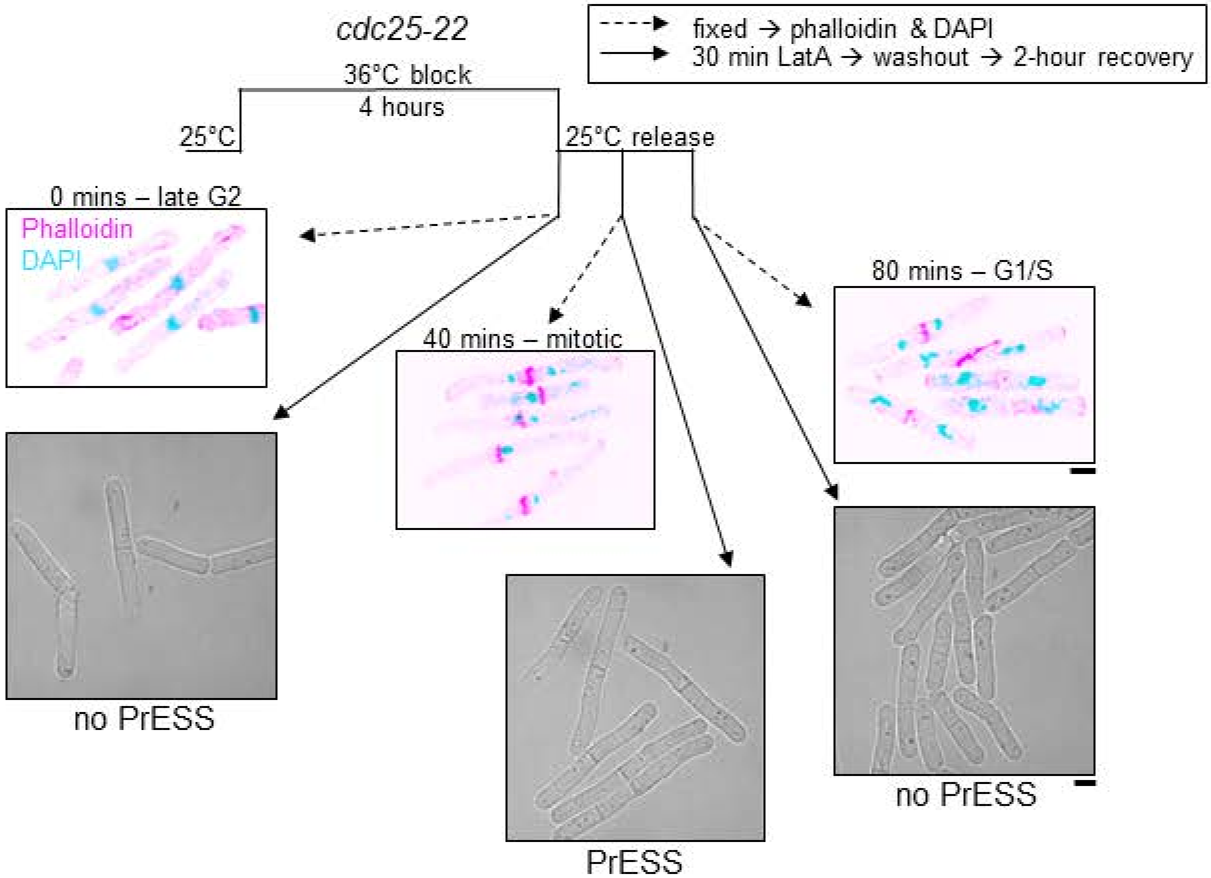


**Supplemental Figure S2:**
**A)** Table showing the frequency of the PrESS phenotype when cells are treated in different cell cycle stages. **B)** *cdc25-22* cells were synchronized by cell cycle block and release. During the indicated cell cycle stages occurring 0, 40, and 80 minutes after release, a fraction of cells was fixed and stained with phalloidin (magenta) and DAPI (cyan), to mark the actomyosin ring and nucleus respectively, while the rest were treated for 30 minutes with LatA and then washed. Cells recovering from LatA treatment in G2 or G1/S does not yield PrESS cells while those undergoing mitosis exclusively yield PrESS cells. Scale bars, 5μm.

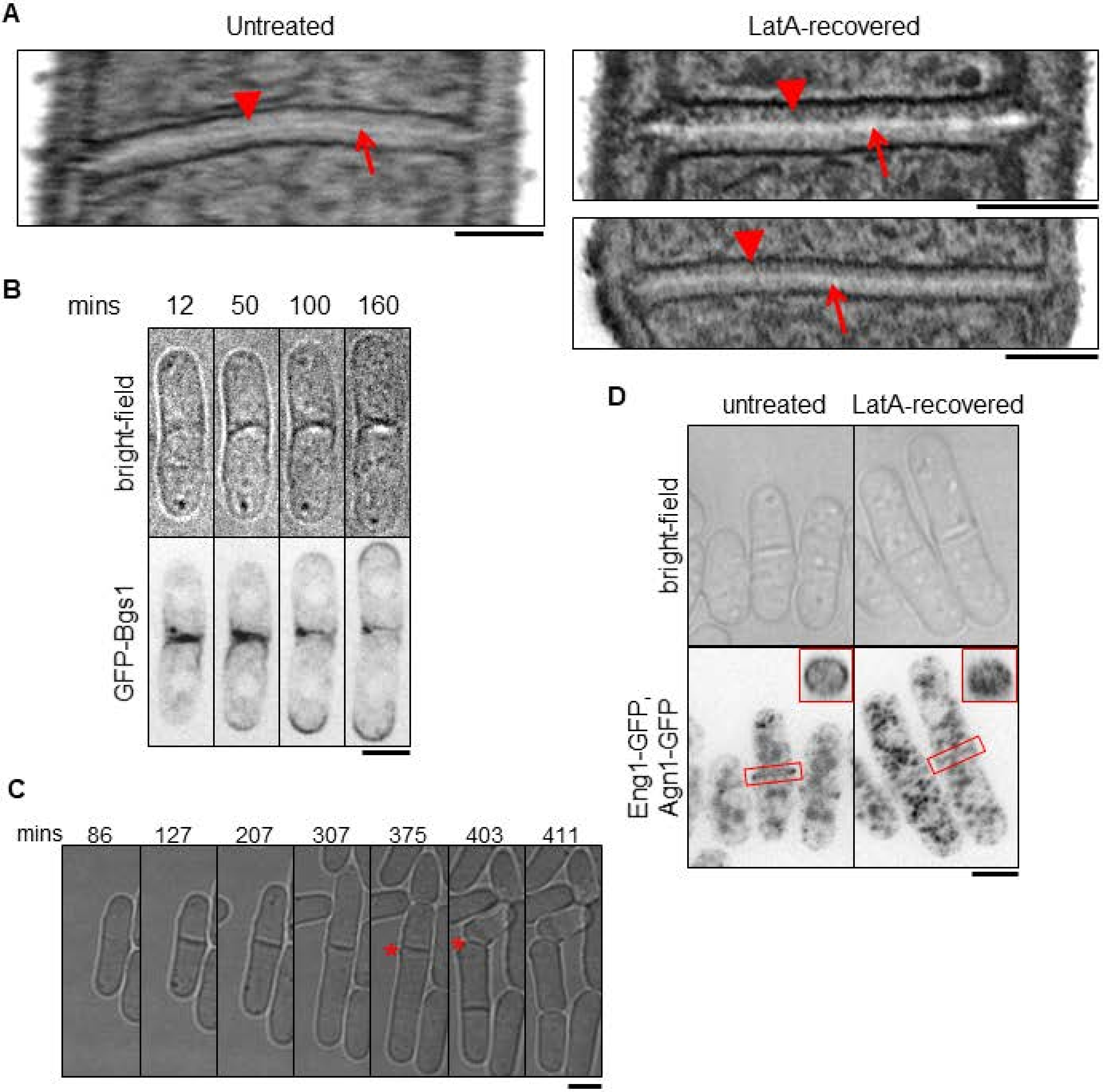


**Supplemental Figure S3**

**A)** Electron micrographs of wild-type septa; untreated (left) and LatA-treated after 2.5-hour recovery (right). Arrows indicate primary septum, while arrowheads indicate secondary septum. Scale bars, 500nm. **B)** Montage of a time lapse of GFP-Bgs1 localization in a PrESS cell. Minutes are since LatA washout. Scale bar, 5µm **C)** The original septum of a PrESS cell sometimes separates (asterisk) albeit after a delay. Minutes are since LatA washout. **D**) Septum-digesting enzymes, Eng1-GFP and Agn1-GFP, in control cells show cortical distribution at the septum, effectuating a ring, while in PrESS cells their localization appears as a disc all over the septum barrier. The LatA-treated cells shown were imaged 3 hours after LatA washout. 3D-reconstructed membrane barriers are shown in the insets. Scale bars, 5µm.

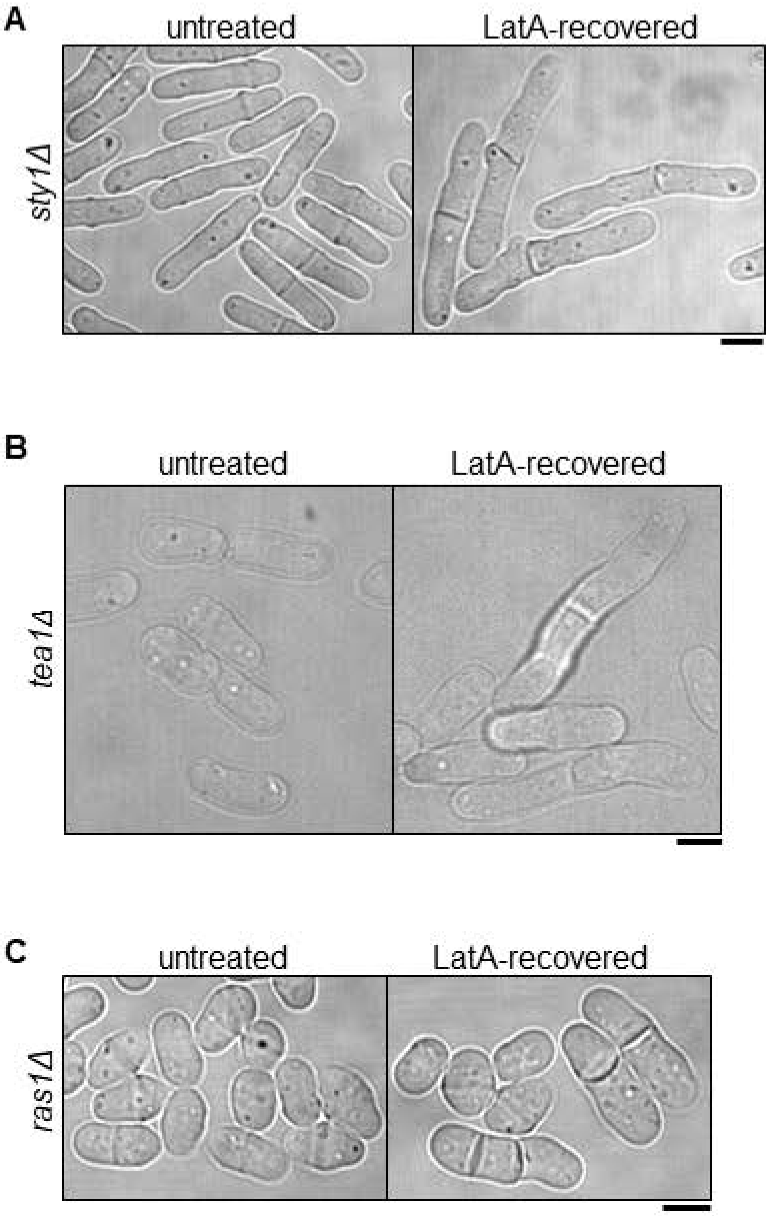


**Supplemental Figure S4**

The PrESS phenotype was observed in mutants of the MAP kinase *sty1Δ* **(A)**, GTPase *ras1Δ* **(B)** and *tea1Δ* **(C)**. Scale bars, 5µm.

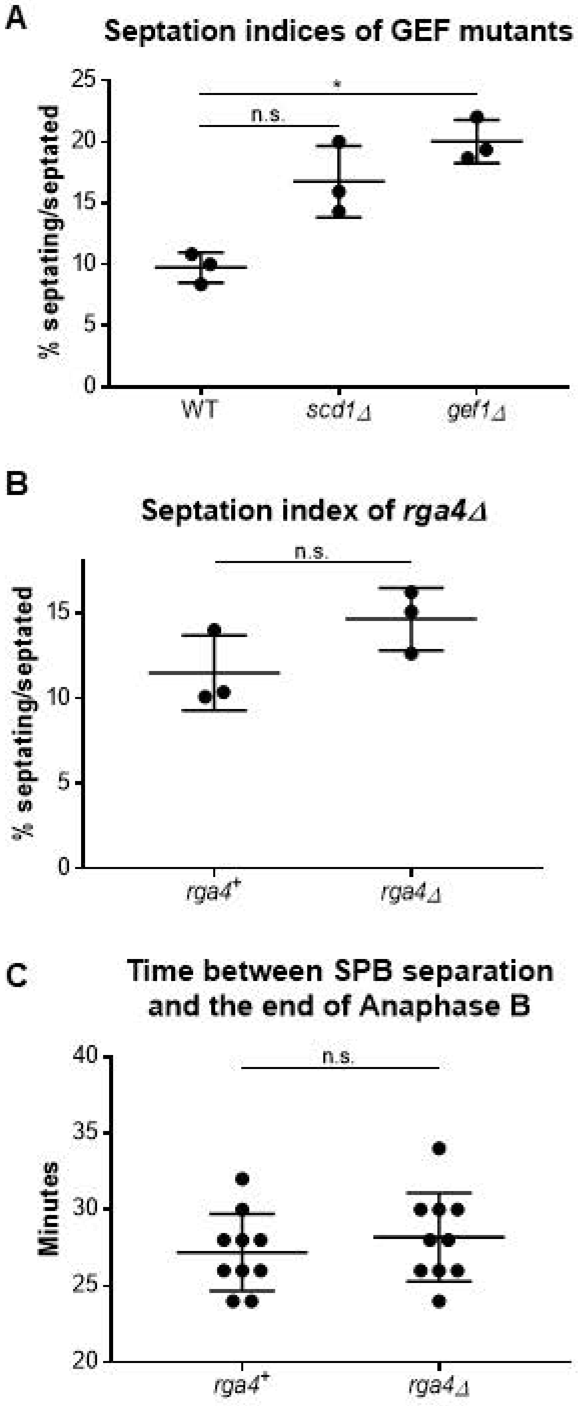


**Supplemental Figure S5**

**A)** Septation indices of the Cdc42 GEF mutants *scd1Δ* and *gef1Δ*. *gef1Δ* shows significantly more septated cells than the wild type ($p = 0.0396$), while *scd1Δ* does not ($p = 0.1017$). **B)** Septation index of the Cdc42 GAP mutant *rga4Δ*, which was similar to wild type ($p = 0.1277$). **C)** Quantification of the time between spindle pole body separation (completion of spindle formation) and the end of Anaphase B. We find no significant difference between *rga4*$^{+}$ and *rga4Δ* cells. Statistical tests used for A is Ordinary one-way ANOVA with Tukey's multiple comparisons and for B and C is Student's t-test.

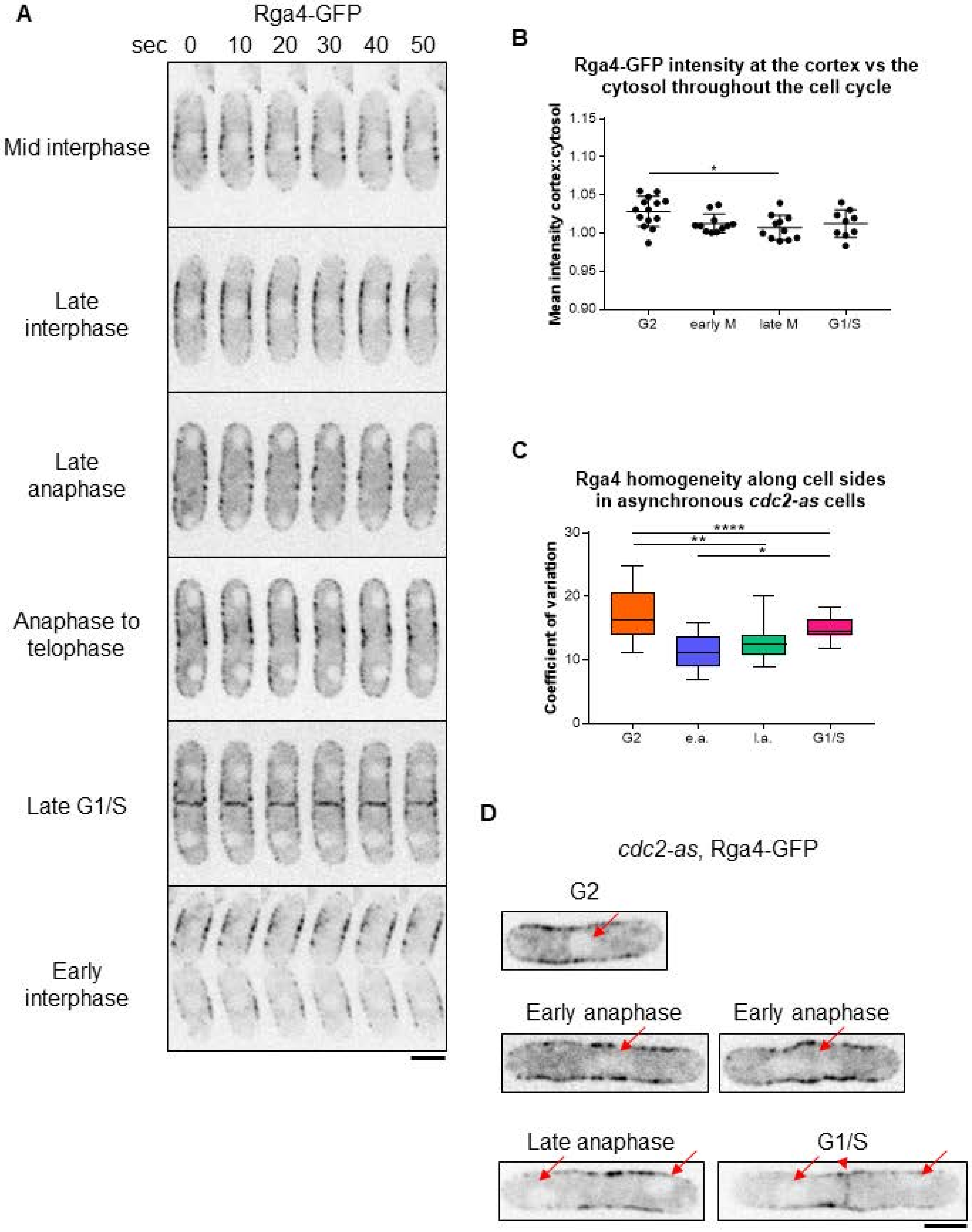

A
Rga4-GFP
sec 0 10 20 30 40 50
Mid interphase
Late interphase
Late anaphase
Anaphase to telophase
Late G1/S
Early interphase
B
Rga4-GFP intensity at the cortex vs the cytosol throughout the cell cycle
Mean intensity cortex:cytosol
1.15
1.10
1.05
1.00
0.95
0.90
G2
early M
late M
G1/S
*
C
Rga4 homogeneity along cell sides in asynchronous cdc2-as cells
Coefficient of variation
30
20
10
0
G2
e.a.
l.a.
G1/S
****
**
*
D
cdc2-as, Rga4-GFP
G2
Early anaphase
Early anaphase
Late anaphase
G1/S

**Supplemental Figure S6**

**A)** Montages of the same cells in Figure 6B. Images are from the first minute of the time-lapse with an interval of 10 seconds. **B)** Quantification of Rga4-GFP intensity throughout the cell cycle. The values are ratios of the intensity at the cortex to the intensity in the cytosol. The intensity of Rga4-GFP is lower in the cytoplasm during G2 than late mitosis ($p = 0.0189$). **C)** Quantification of the homogeneity of Rga4-GFP in asynchronous *cdc2-as* cells without inhibitor treatment. e.a. = early anaphase; l.a. = late anaphase. Ordinary one-way ANOVA with Tukey's multiple comparisons. **D)** Sample Rga4-GFP expressing *cdc2-as* cells demonstrating how cells in different cell cycle stages were selected. Arrows show a single nucleus in G2, an elongated nucleus in early anaphase, a binuclear cell with no signal at the division site for late anaphase, and a binuclear cell with signal at the division site (arrowhead) for post-anaphase cells. Scale bars, 5μm.

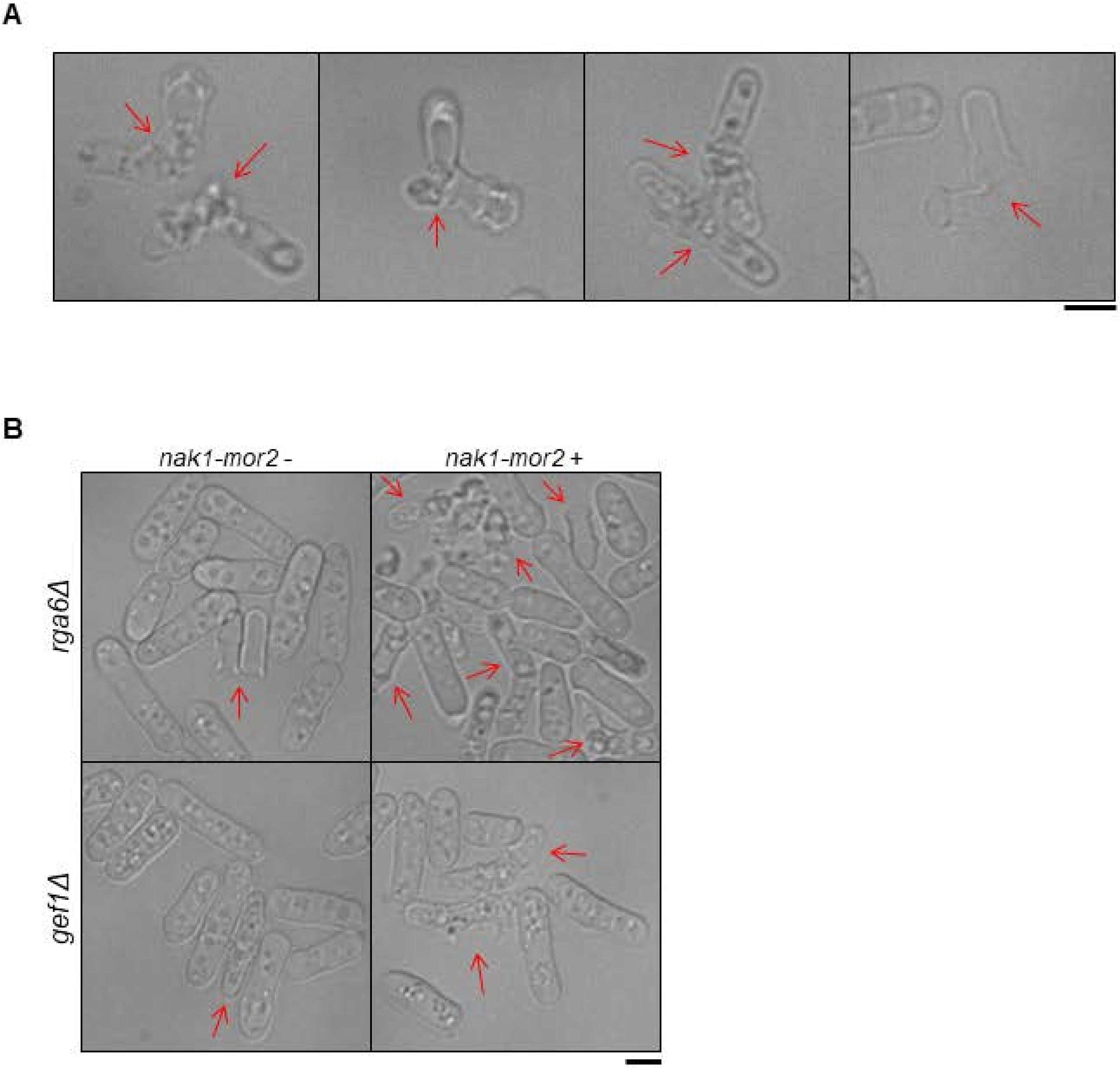


**Supplemental Figure S7**

**A)** Examples of lysed cells (arrows) in wild type cells expressing *nak1-mor2* (-thiamine). Cells mostly lyse at the division site. **B)** *gef1Δ* and *rga6Δ* cells under conditions where they repress *nak1-mor2* or express *nak1-mor2*. Lysed cells are shown with arrows. Scale bars, 5μm.
