## Supplemental Text for "Cdc42 reactivation at growth sites is regulated by local cell-cycle-dependent loss of its GAP Rga4"

**Table S1: Strains list**

| Strain | Genotype | Source |
|---|---|---|
| PN567 | *h+ ade6-704 leu1-32 ura4-d18* | Paul Nurse |
| FV467 | *h+ tea1Δ::ura4+ ura4-D18 ade6-M210* | (Verde et al., 1995) |
| FV513 | *h- rga4Δ::ura4+ ade6-704 leu1-32 ura4-d18* | (Das et al., 2007) |
| FV784 | *h+ rga4-gfp::KanMX ade6-704 ura4-D18 leu1-32* | (Das et al., 2007) |
| JM1745 | *h- sty1Δ:ura4+ leu1-32 ura4-D18* | (Millar et al., 1995) |
| JX125 | *h90 Δscd1::ura4+ ade6 leu1-32 ura4-d18 h210* | (Hirota et al., 2003) |
| JZ521 | *h90 ras1Δ::ura4+ ade6-M210 leu1-32 ura4-D18* | (Hakuno et al., 1996) |
| PPG2601 | *h+ Δgef1::ura4+ ura4-D18 leu1-32* | (Coll et al., 2003) |
| PPG3750 | *h- bgs1Δ::ura4 Pbgs1::GFP-bgs1:leu1 leu1-32 ura4-D18 his3-D1 h-* | (Cortes et al., 2002) |
| PPG5668 | *h+ scd2-GFP-kanMX6+ leu1-32 ura4-d18* | (Rincon et al., 2007) |
| WF206 | *h- cdc25-22 ade6-704 leu1-32 ura4-D18* | (Russell and Nurse, 1986) |
| YSM740 | *h+ myo52-tdTomato-NATrMX ade6-M216 leu1-32 ura4-D18* | (Martin et al., 2007) |
| YSM947 | *h+ scd1-3GFP-kanMX+ ade6-m216 leu1-32 ura4-d18* | (Bendezu and Martin, 2013) |
| YMD317 | *CRIB-GFP-ura4+ rlc1-Tomato-NATr sad1-mCherry:kanMX ade6-M21X leu1-32 ura4-D18 his7+* | Lab stock |
| YMD546 | *bgs1Δ::ura4 Pbgs1::GFP-bgs1:leu1+ Rlc1-tdTomato-NATr Sad1-mCherry:kanMX leu1-32 ura4-D18* | (Wei et al., 2016) |
| YMD772 | *rga4-GFP-KanMX rlc1-tdTomato-NATr sad1-mCherry:kanR ade6-M216 leu1-32 ura4-D18 his7+* | Lab stock |
| YMD824 | *eng1-GFP-KanMX agn1-GFP-KanMX leu1-32 ura4-D18* | This study |
| YMD1200 | *h- [pREP41X-Nak1-Mor2] ade6-704 leu1-32 ura4-d18* | This study |

| | | |
|---|---|---|
| YMD1253 | *[pREP41X] CRIB-GFP-ura4+ ade6-704 leu1-32 ura4-d18* | This study |
| YMD1255 | *[pREP41X-Nak1-Mor2] CRIB-GFP-ura4+ ade6-704 leu1-32 ura4-d18* | This study |
| YMD1509 | *h90 cdc2-as-bsd rga4-GFP::KanMX leu1-32 ura4-D18 ade6-M216* | This study |
| YMD1512 | *h- rga4Δ::ura4+ [pREP41X-Nak1-Mor2] ade6-704 leu1-32 ura4-d18* | This study |
| YMD1735 | *h- rga6Δ::ura4+ [pREP41X-Nak1-Mor2] ade6-704 leu1-32 ura4-d18* | This study |
| YMD1737 | *h- gef1Δ::ura4+ [pREP41X-Nak1-Mor2] ade6-704 leu1-32 ura4-d18* | This study |
| YMD1741 | *h- rga6Δ::ura4+ [pREP41X] ade6-704 leu1-32 ura4-d18* | This study |
| YMD1743 | *h- gef1Δ::ura4+ [pREP41X] ade6-704 leu1-32 ura4-d18* | This study |
| YMD1831 | *h- rga4Δ::ura4+ [pREP41X-Nak1-Mor2] CRIB-GFP-ura4+ ade6-704 leu1-32 ura4-d18* | This study |
| YMD1835 | *h- rga4Δ::ura4+ [pREP41X] CRIB-GFP-ura4+ ade6-704 leu1-32 ura4-d18* | This study |

**Table S2: Candidate screen of mutants to identify changes in the PrESS phenotype.** All mutants listed here showed the PrESS phenotype during LatA recovery unless marked with an asterisk.

| **Protein** | **Mutant** | **Role** |
|---|---|---|
| Cdc16 | *cdc16-116****** | SIN inhibitor |
| Scd1 | *scd1Δ* | Cdc42 GEF |
| Gef1 | *gef1Δ* | Cdc42 GEF |
| Gef1 | *gef1S112A* | Cdc42 GEF |
| Rga4 | *rga4Δ* | Cdc42 GAP |
| Rga6 | *rga6Δ* | Cdc42 GAP |
| Scd2 | *scd2Δ* | Scaffold for Scd1 |
| Rdi1 | *rdi1Δ* | Cdc42 GDI |
| Ras1 | *ras1Δ* | RAS GTPase |
| Tea1 | *tea1Δ* | Polarity marker |
| Tea4 | *tea4Δ* | Polarity marker |
| For3 | *for3Δ* | Actin regulator |
| Myo1 | *myo1Δ* | Type I myosin |
| Sec8 | *sec8-1* | Exocyst component |
| Exo70 | *exo70Δ* | Exocyst component |
| Sty1 | *sty1Δ* | MAP kinase |
| Orb2 / Shk1 / Pak1 | *orb2-34* | PAK kinase |
| Plo1 | *plo1.as8* | Polo kinase |
| Cdc10 | *cdc10-129* | Cell cycle regulator |

### Supplementary movies

**Supplementary movie S1: PrESS Cells.** Cells treated with 10µm LatA for 30 minutes and washed and allowed to recover. Cells initiate growth at the ends before the septum matures and fail to separate.

**Supplementary movie S2:** Time-lapse movie of a cell in mitosis undergoing LatA treatment, then washout after 30 mins. This cells recovers and initiates growth (asterisk) soon after ring closure. Upper panel is brightfield imaging, lower panel is Rlc1-TdTomato to mark the actomyosin ring and Sad1-mCherry to mark the spindle pole bodies.

**Supplementary movie S3:Rga4-GFP dynamics in different cell-cycle stages.** *rga4-GFP*-expressing cells were imaged every 10 secs for 5 mins. **A)** A cell in mid G2 phase. **B)** A cell in late G2 phase. **C)** A cell in late anaphase. **D)** A cell in telophase. **E)** A cell in late G1/S phase. **F)** A cell in early G2 phase.
